# Functional and compositional diversity peak at intermediate fire frequencies when modeling the plant-fire feedback

**DOI:** 10.1101/2025.06.10.656287

**Authors:** Matilde Torrassa, Gabriele Vissio, Rubén Díaz Sierra, Marta Magnani, Maarten B. Eppinga, Mara Baudena

## Abstract

Fires are generally considered to promote biodiversity, although the exact relationship is unclear, because it can be affected by several factors, including fire regime and ecosystem type. Given the ongoing global change, a better understanding of this connection is needed to assess the extent to which projected increases in fire frequency may affect current biodiversity trends. A major challenge lies in vegetation-fire feedback, which often mediates changes in fire regimes. To shed light on the role of fires in promoting or limiting biodiversity, we studied the compositional and functional diversity of simulated plant communities along a gradient of fire frequencies. We extended an existing model to include a large number of species. The model reproduces plant successional dynamics and is parameterized to represent Boreal and Mediterranean communities. Fire events are stochastic, with frequencies that depend on community flammability, and plants have different fire responses, thus creating a vegetation-fire feedback. For both ecosystems, we found that fires generally had a positive effect on both compositional and functional diversity. Furthermore, in most cases, both peaked at intermediate fire frequencies. Interestingly, compositional and functional diversity were correlated but did not reach their maximum values in the same communities. This seemingly underlines that a certain degree of functional similarity may be necessary to achieve maximum species richness. These results stem from the vegetation-fire feedback, highlighting its importance for predicting ecosystem responses to global change, including biodiversity losses and wildfire regime shifts.

## Introduction

Fires play a key role in the Earth system, as they occur in most ecosystems that support sufficient biomass to burn (Archibald et al., 2013). Over millennia, fires have been crucial in shaping the composition and distribution of biomes (Bond and Keeley, 2005) since they first appeared, concomitant with the origin of terrestrial plants (Pausas and Keeley, 2009). Fire occurrence is strongly controlled by climate, and requires specific weather conditions (Boer et al., 2017). At the same time, wildfires are also dependent on vegetation, which constitutes the main fuel for fires (Bowman et al., 2020).

In fire-prone ecosystems, vegetation influences the local fire behavior through its flammability, which arises from key functional traits of plants relating to, e.g., the quantity, quality, moisture content, and aeration of plant biomass (e.g. Pausas and Keeley, 2021; Simpson et al., 2022). Plants have also adapted to survive and thrive in fire-prone ecosystems, evolving strategies that enhance the survival probability of populations or individuals. Combustion-stimulated germination or post-fire seed release in crown systems (i.e., serotiny) grant population persistence, whereas plant individual persistence is granted by the ability to resprout after fires. Resprouting forms part of a set of plant strategies connected to protect vital tissues above and below-ground (Clarke et al., 2013). Together, these strategies are the so-called fire responses (Pausas, 2019). The combination of plant traits related to flammability and to fire responses can lead to the emergence of a positive vegetation-fire feedback, playing a key role in (sub-)tropical, temperate, and boreal ecosystems around the world (Staver et al., 2011; Stephens et al., 2014). A positive feedback can emerge in the presence of fire-prone vegetation that stimulates fires and also has strong post-fire recovery: more fires enhance plant growth, in turn leading to a larger number of fires. This is the case, for example, in tropical grasslands and savannas, dominated by flammable and fire-stimulated C_4_-grasses (Bond, 2008). The same positive feedback can also sustain ecosystems where fires rarely occur, such as some tropical forests, where vegetation limits fire spread by creating a humid environment and reducing wind speeds, while at the same time not having any specific response to fire (e.g. Charles-Dominique et al., 2017). Plant community composition thus responds to and influences the frequency, intensity, and extent of fire events, that is the fire regime (Archibald et al., 2013). The prevailing fire regime is then the result of climatic and meteorological drivers, mediated by these fire-vegetation feedbacks (Archibald et al., 2018).

Fire has been identified as a critical driver of biodiversity and ecosystem functioning in fire-prone ecosystems (Bond and Parr, 2010; He et al., 2019; Kelly et al., 2020; Pausas and Keeley, 2019). Understanding the connection between wildfires and biodiversity in different fire-prone systems around the world is an important challenge given the ongoing biodiversity crisis and the new era of fire regime, with increasingly intense and extensive fires on the one hand, and increased fire suppression on the other (Cunningham et al., 2024; Kelly et al., 2020). Generally, fires are thought to promote turnover and diversification of plant populations, and can promote the coexistence of plant species (Brooks et al., 2004; He et al., 2019; Pausas and Ribeiro, 2017, and references therein): They reduce the dominance of the most competitive plant species and allow different species to dominate at different stages of post-fire succession. However, extreme conditions, e.g., very frequent or very infrequent fires, may reduce the overall biodiversity. Specifically, theory predicts that very frequent fires seemingly tend to exclude all plant species that have no response to fires, but also, for example, those with seedbanks that require longer periods to reach maturity, leading to population declines and even local extinctions (e.g. Bowman et al., 2013; Enright et al., 2015). Conversely, when fires are absent for decades or centuries, many shade-intolerant or fire-dependent species (e.g., conifers with serotinous cones) will have been excluded by stronger competitors for light and other limiting resources, or disappeared due to a lack of viable offspring (He et al., 2019, and references therein). Given these premises, diversity is expected to peak somewhere in the middle between these two extreme scenarios, where fire-adapted and unadapted plant species can coexist. This is similar to what is postulated by the *Intermediate Disturbance Hypothesis* (IDH) (Connell, 1978; Graham and Duda, 2011; Huston, 1979; Willig and Presley, 2018), even though the IDH does not consider the fact that fire is an endogenous element in plant ecosystems (Bond and Keeley, 2005; Kelly et al., 2020). A hump-shaped relationship between plant compositional diversity and fire frequency has been supported in several ecosystems (Dee and Menges, 2014; Gordijn et al., 2018; Pellegrini et al., 2023). In some studies, however, species richness monotonically decreases with increasing fire frequency, especially among woody species. In fact, it has been argued that such discrepancies might be due to the difficulties in sampling a sufficiently large set of fire frequencies, or in sampling systems with species turnover along the fire frequency gradient, rather than capturing responses of a fixed species pool (He et al., 2019, and references therein). In a set of model experiments where fires were imposed as exogenous disturbances, a multimodal diversity-disturbance relationship emerged (Liao et al., 2022). Therefore, the consensus on the hump-shaped relationship between species richness and diversity on one hand, and fire frequency on the other, is not fully established. Furthermore, while it is likely that different fire regimes are related to changes in plant functional traits (e.g. de L. Dantas et al., 2013; Pacuk et al., 2025), functional diversity changes across different ecosystems and fire regimes show contrasting patterns and have not yet been investigated comprehensively (He et al., 2019; Meza et al., 2023; Schaffhauser et al., 2012).

Here we set out to study how compositional and functional plant diversity vary along gradients of fire frequency, across two important and iconic fire-prone systems, namely Mediterranean forests and shrublands, and Boreal forests, where fires are expected to increase in the future (Seidl et al., 2020; Shen et al., 2025; Turco et al., 2018). To do so, we extend an existing theoretical model of plant and fire dynamics (Magnani et al., 2023), to include a large number of species. The model represents plant succession (following Tilman, 1994), species-specific fire responses, and fires as stochastic events whose occurrence depends on the flammability of plant communities. Thus, the model captures the existence of feedback between plant community composition and fire frequency. We parameterize the model for real communities in the Mediterranean and Boreal ecosystems. Our analysis consists of three steps, answering the following research questions: i) How do fires impact compositional and functional diversity in Mediterranean and Boreal ecosystems, as compared to systems without fires? ii) How do both types of diversity depend on the fire frequencies that emerge within the fire ecosystems? iii) To what extent is a higher number of coexisting plant species associated with a larger range of competitive and fire response strategies? Our results show that fires promote biodiversity, as expected; furthermore, diversity indicators peak at intermediate fire frequency. The two types of diversity are not always strongly correlated within communities.

## Methods

### Competition model with stochastic fires

In order to explore fire-driven biodiversity patterns in a general framework, we developed a conceptual model describing the dynamics of fire-prone plant communities, based on and extended from the work in Magnani et al. (2023). The model includes competitive plant dynamics during fire-free periods, stochastic fires, and plant fire responses. We extended the original approach of Magnani et al. (2023), which represented three species, to represent a large number (*N*) of plant species.

The model represents the competition-colonization dynamics of plant species, following the classical approach of Tilman (1994). This is a proximal representation of plant competition for a single resource (mostly light in this and previous works (Baudena et al., 2020; Magnani et al., 2023)). Each species has a fixed position in the competition hierarchy, from the strongest (*i* = 1) to the weakest (*i* = *N*). The governing equation of the model for the plant cover *b_i_*(*t*) of species *i* is (analogously to Tilman, 1994):

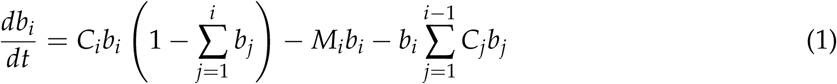

In this representation, each plant species can colonize both the empty space and the space occupied by inferior competitors. *C_i_* is the maximum per capita colonization rate, which is multiplied by the population size *b_i_* to obtain the fraction of space that species *i* would colonize per time unit in the absence of other species. This rate is then reduced by the crowding factor 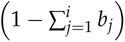, which quantifies the spatial restrictions to colonization due to the presence of superior competitors or of species *i* itself. *M_i_* is the mortality rate, whereas the last term on the right-hand side of eq. 1 represents the superior species’ skill to invade and colonize species *i* space, and can therefore be defined as a displacement term. In the absence of hierarchically superior species, persistence of a species requires satisfying a simple condition, *C_i_* − *M_i_ >* 0. This condition of existence is necessary for each species, but for species lower in the hierarchy (*i >* 1) it is not sufficient to ensure survival, due to the displacement term.

Fire events are modeled as instantaneous, stochastic events, with a probability of occurrence that is exponentially distributed. The average fire return time (i.e., the mean time interval between two consecutive fire events) is inversely proportional to the flammable plants vegetation cover (Baudena et al., 2020; Magnani et al., 2023):

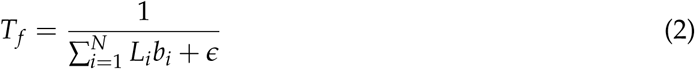

where *L_i_* is the flammability of the species *i*, and the term *ϵ* = 10^−3^*yr*^−1^ ensures the convergence of fire return time when the total plant cover is null. The stochasticity of the fire signal is simulated using a nonstationary Poisson process (Sigman, 2007) with rate *λ* = 1/*T_f_*. The interarrival time of fires thus follows an exponential distribution, with a mean interarrival time depending on the vegetation cover and community composition in the preceding time-step. Note that the mean interarrival time responds instantaneously to changes in community composition, meaning that there is no delay considered within the fire-vegetation feedback. Furthermore, we imposed a realistic minimum fire return time of 2 years (following Baudena et al., 2020; Magnani et al., 2023).

Plant response to fire is represented by the parameter 0 ≤ *R_i_* ≤ 1. When a fire occurs, the cover of each species is reduced to a specific fraction determined by its fire response,

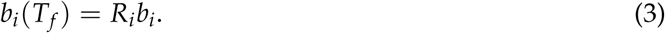

We follow here the formulation introduced by Magnani et al. (2023), where the fire response *R_i_* is a parameter that represents the different plant fire strategies at individual and population level. In this approach, and in agreement with Pausas and Lavorel (2003), the different fire responses are ranked from no response for very low *R_i_* values, to intermediate values for responses at population level (e.g., re-seeding), to high values when responses at individual level (e.g., resprouting) are present.

We investigated the relationship between diversity and fire return time in two fire-prone systems: Mediterranean forests and shrublands, and Boreal forests. In Mediterranean-type regions, the prevailing vegetation is composed of evergreen sclerophyllous-leaved shrublands, semi-deciduous scrub, and woodlands, whose high flammability leads to frequent, high-intensity summer fires (Keeley, 2012). These regions are often densely inhabited, and fire ignition is usually attributable to anthropogenic causes (Ganteaume et al., 2013). Boreal ecosystems are typically dominated by conifers with a ground layer vegetation of shrubs, low herbs, and moss. Aside from anthropogenic activity, fire ignition is often due to lightning, and fire hazard is spread over all seasons (Rowe and Scotter, 1973).

In an effort to represent realistic plant communities, for the parameters related to succession (*C_i_* and *M_i_* in eq. 1), we used values from Baudena et al. (2020) and Magnani et al. (2023) (see tables A1-A2). In these studies, these parameters were derived from literature and expert estimation, or calibrated from field data. For the Mediterranean case we obtained values from Baudena et al. (2020) for six species, while for the Boreal case three plant functional types from Magnani et al. (2023). To extend to a larger number of species, we assumed that the *C_i_* and *M_i_* values were related to the species’ position in the hierarchy *i*: we thus performed linear regressions in which the colonization and mortality rates were the response variables and the position in the hierarchy was the predictor variable. We then assumed that these relationships were generalizable to all our *N* species, and thus we generated their parameters *C_i_* and *M_i_* stochastically from the fitted regression models:

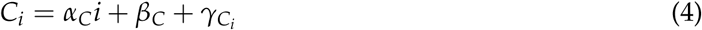

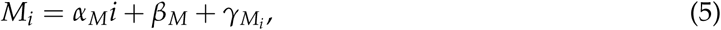

where *α*s and *β*s are, respectively, the slopes and the intercepts of the regressions in fig. A1, and *γ*s are zero-mean gaussian noise terms, whose standard deviations are also determined from the standard deviation of the data around the linear regression (see table A3 and the vertical bars in fig. A1). As a further requirement, we imposed that each generated species must satisfy the condition *C_i_* > *M_i_ >* 0, i.e., it must be able to potentially persist in isolation. Moreover, as in the present paper we set *N* to different values, we rescaled the hierarchy so that the first and *N*-th species represent the boundaries of the parameter range provided by the empirical data underlying the regression models depicted in fig. A1. From the linear fits, the colonization rates are inversely related to the position in the hierarchy. This is reasonable since generally plants with better access to light are also slower in growing and colonizing than the less competitive ones — that is, trees generally grow slower than shrubs and grasses. Interestingly, in the two ecosystems the regression of mortality rate as a function of hierarchy presents opposite trends, thus ensuring more generality to communities generation and, consequently, to the model results. We also generated a second set of simulations to include annual plants in the Mediterranean (see Supporting information for details). We did this to overcome the fact that our parameterization, following Baudena et al. (2020), encompassed growth rates and mortalities that represented a variety of life forms, spanning from perennial grasses to late successional trees, but excluded annuals, which are very relevant in Mediterranean ecosystems (Poppenwimer et al., 2023).

Finally, to explore a comprehensive variety of fire strategies, leading to a wide assortment of fire return times, we did not impose fixed values on the ecosystem’s fire-responses and flamma-bilities — which lead to the fire-vegetation feedback. Thus, a variety of fire trait combinations was simulated. Fire response *R_i_*, describing the fraction of plant cover persisting after a fire event, due to either plant resistance or recovery, was sampled from a uniform distribution between 0.001 and 1. Flammability is defined as *L_i_*= 10^−^*^θ^ y*^−1^, where *θ* is a random number uniformly distributed between *log*_10_(2) and *log*_10_(500) — thus, 0.002 ≤ *L_i_* ≤ 0.5 yr^−1^. In this way, we represented an average fire return time varying between 2 and 500 years, which includes the typical fire frequencies of the ecosystems studied (Harrison et al., 2021; Henne et al., 2013; Nasi et al., 2002).

### Simulations

We generated and simulated 1000 communities for each of the following cases: Mediterranean communities with i) *N* = 10 initial species (hereafter Med-10) and ii) with *N* = 50 initial species (hereafter Med-50); and iii) Boreal type communities with *N* = 10 initial species (hereafter Bor-10). The limited number of initial species in the Boreal communities reflects the smaller species pool and lower biodiversity of the Boreal ecosystems compared to those of the Mediterranean. In fact, among these three types of communities, Med-50 and Bor-10 can be considered the most realistic, with Med-10 providing a reference to study the separate effects of initial species richness (through comparison with Med-50) and the effect of ecosystem type (through comparison with Bor-10).

#### Initial conditions

Every community, initially composed of *N* species randomly generated as explained in section “Competition model with stochastic fires”, underwent *N* + 2 runs, each with different initial conditions, determined as follows:

- 1 initial condition corresponding to small and equal vegetation cover for all species:

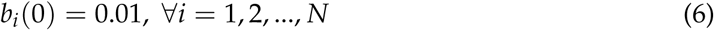
- 1 initial condition corresponding to equal vegetation cover for all species, with larger values than in the previous case, given by:

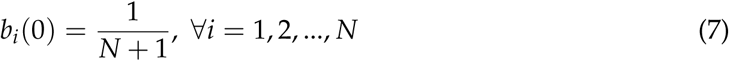
- *N* initial conditions with high vegetation cover for a single species and low for all the others, leaving an initial portion of empty space equal to 0.1:

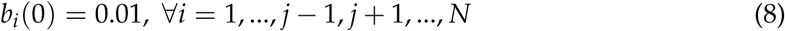

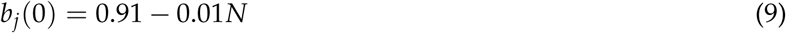

Such initial conditions are designed to meet the requirement of total vegetation cover not exceeding the unity (Tilman, 1994). In the second initial condition, we set the denominator to *N* + 1 in order to provide some free space for the species to grow, as it is known that the sum of the plant equilibria in Tillman’s model is always below 1.

#### Numerical integrations

The model has been coded in Fortran; some results were also reproduced with Python code (see section “Data and Code Availability”). All the numerical integrations presented below are performed using a Runge-Kutta 4 method, with runs of 10^4^ *y* and time steps of 1 *day*.

The different set of initial conditions brings the total number of simulations to 12,000 for Med-10 and Bor-10 and to 52,000 for Med-50. Such large number of runs, together with their duration, allows for a proper statistical representation of these stochastic systems (Baudena et al., 2020; Magnani et al., 2023).

### Analyses

We analyzed the time series obtained in the second half of each run, i.e., 5,000 years, considering the first half as spin-up time needed to reach a statistically steady state. Over the latter interval, we averaged each species’ cover, acquiring a set of average cover values ⟨*b_i_*⟩, which defined the state achieved by a community starting with a specific set of initial conditions. Furthermore, for every run we averaged the system’s fire return time, obtaining a value ⟨*T_f_* ⟩ corresponding to each set of ⟨*b_i_*⟩s. As shown in Magnani et al. (2023), running this model (with *N* = 3) multiple times can result in different final communities even when using species with the same set of parameters, due to different initial conditions and/or to the stochastic signal underlying fire occurrence. Thus, we expect multiple stable states are likely to exist also in the model with *N >* 3, and we chose a set of initial conditions and a large enough number of simulations to capture such alternative ecosystem states, as they possibly enlarge and enrich the diversity of the final communities represented. Yet, we deem that a thorough identification of all possible ecosystem states is not necessary to reach the aims of this paper. Simply, we avoided replicating states by discarding duplicate communities resulting from the same set of simulations. To deter-mine whether community ⟨*b*^′^⟩ was a duplicate of community ⟨*b*⟩, we considered their absolute difference:

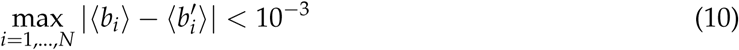

In the following, we introduce how we estimated the biodiversity of the simulated ecosystems quantitatively, using four indicators that account for both compositional and functional diversity, and how we answered the three research questions by analyzing their relationships with fire return time.

#### Compositional diversity indicators

We estimated ecosystems’ biodiversity using a set of biodiversity indicators based on the *composition* of the plant communities (Noss, 1990), namely Species Richness and the Inverse Simpson Index. They are defined and estimated as follows:

- *Species Richness* (*S*) is the number of coexisting species in the community (considering as minimal threshold for the presence of each species an average vegetation cover ⟨*b*⟩ ≥ 10^−3^).
- *Inverse Simpson Index* (*I*) evaluates the inverse probability of different species to exist within the community (Simpson, 1949) and is defined as:

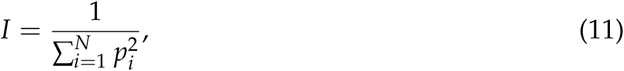

where *p_i_*is the relative abundance of species *i*, calculated from cover as:

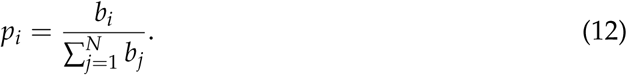

Note that, in each summation, only *S* species survive among the *N* initial species. Therefore, *N* − *S* terms in the denominator of eq. 12 are null.

#### Functional diversity indicators

Another important biodiversity attribute is the *function*, which involves ecological and evolutionary processes, including gene flow, disturbances, and resource utilization and cycling (Noss, 1990). We implemented functional diversity indices as represented in our model by the diversity of the plant species parameters in the final community assembly. To do so, we used the concept of *hypervolumes*, that is the volume in the trait (or parameter, in our case) space of the surviving species (Blonder et al., 2014; Mammola and Cardoso, 2020), and estimated for each community assembly two aspects of functional diversity (Mason et al., 2005): Functional Richness and Divergence, as described in the following.

We considered the parameter space composed of the colonization rate *C*, the fire response *R*, and the flammability *L*. We excluded the competition index *i* and the mortality rate *M* as we imposed a relationship between them and *C*. For the hypervolume estimation, the composition of each community was transformed into an input dataset, weighted using the species cover: each plant parameter set was repeated in the dataset ⟨*b*⟩ · 10^3^ times (rounded), consistently with the threshold set to consider species presence in the resulting community. This had the effect of weighting the parameters according to the species cover.

To ensure each axis had comparable units (Blonder et al., 2014), the parameters were rescaled using a range normalization: with *t* being the original value of a parameter, *t_min_* and *t_max_* the minimum and maximum value of that parameter, the parameter was normalized as *t*^′^ = (*t* − *t_min_*)/(*t_max_* − *t_min_*). Fire response *R* and flammability *L* range values were set as equal for all communities simulated in the two ecosystem types, given that their range values were fixed and common to both cases. Conversely, the colonization rate *C* normalization was based on the resulting parameter range values for each ecosystem type, depending on the communities’ generation values.

The hypervolume of each community was then computed as a rescaled sample inferred from a Gaussian (KernelDensity function of the sklearn.neighbors Python library) of the input community datasets. The representation of an estimated hypervolume is reported in fig. B1 as an example. The calculation method implemented for the hypervolume estimation is a simplification of the original proposed by Blonder et al. (2014), which was developed to model high-dimensional and irregular populations (e.g., with holey distributions), while our modeled communities represent a relatively low-dimensional trait space (3*D*) and consist of sets of 50 unique points at most.

We then estimated the functional richness and divergence metrics from the constructed hypervolumes:

- *Functional Richness* (*F_R_*), which is the total amount of trait space available, was computed as the volume of the Convex Hull derived from the input hypervolume, using the spatial.ConvexHull.volume function of the scipy Python library (Virtanen et al., 2020).
- *Functional Divergence* (*F_D_*), which measures the density of the trait space, was computed as the average distance between the trait space centroid and random points within the boundaries of the hypervolume, using the staila.distance.cdist function from the scipy Python library (Virtanen et al., 2020) to estimate the 3*D* distance.

The functional richness and divergence metrics were constructed based on those proposed by Cardoso et al. (2015) in BAT - Biodiversity Assessment Tools (kernel.alpha and kernel.dispersion functions, respectively). See section “Data and Code Availability” for the full implementation of the code.

Although these indices are known to generally correlate with species richness, this limitation is reduced in systems strongly dependent on initial conditions, i.e., where assembly order can influence the final community composition (Mason et al., 2013). The present model satisfies both these requirements, as shown by Magnani et al. (2023) for the *N* = 3 species version, supporting the implementation of these metrics in our case.

#### Fire-biodiversity analyses

To answer the first research question and investigate the role of fires for community diversity, we analyzed the difference in diversity indicators between communities simulated with and without fires — the latter case being the Tilman (1994) model. Taking the Species Richness difference as an example, we estimated Δ*S* = *S*_fire_ − *S*_no-fire_, with *S*_fire_ being the Species Richness of a community resulting from our model simulation, and *S*_no-fire_ the Species Richness of the community resulting from the model without fires — which we simply calculated analytically as the equilibria of the Tilman (1994) equations for the same initial set of species. The distributions of these biodiversity-indicator differences were tested to assess whether the difference between indicators built on paired ecosystems — i.e., the final communities achieved after running the model with and without fires — was significantly different from zero. Inverse Simpson Index, Functional Richness, and Functional Divergence difference distributions were tested through the Wilcoxon signed-rank test. Since the rank test cannot be applied to discrete distributions, Species Richness difference distributions were tested via the Sign Test. The tests were conducted using respec-tively the scipy.stats.wilcoxon function of Python library SciPy (Virtanen et al., 2020) and the statsmodels.stats.descriptivestats.sign test function of Python library StatsModels (Seabold and Perktold, 2010). Furthermore, we also compared visually for each community the values of the different diversity indicators simulated with and without fires along the fire return time axis resulting from the simulations.

To answer the second question, we analyzed the biodiversity indicators of the resulting communities as a function of their fire return times. We performed quantile regressions to test the significance of the relationship, since the biodiversity indicators are bounded to minimum values (1 for *S* and *I*, 0 for *F_R_* and *F_I_*), and the homogeneity of their variance across the fire return time gradient could not be assumed (Belyea, 2007). We used the 95*^th^*, 90*^th^*, and 85*^th^* quantiles to focus on the trends of the diversity maxima and to estimate the functional relationship near the upper limits of the data distribution (Belyea, 2007, and references therein). For each quantile, we fitted the data with first- to fifth-order polynomial quantile regressions and selected the most performing polynomial regressions based on the Akaike Information Criterion and the Walt Test of significance (Koenker and Portnoy, 1996).

For the third question, we estimated the correlations between compositional and functional diversity indicators to investigate whether the different indices estimated from the same simulated communities resulted in similar patterns. To test for their correlation, we used the Spearman rank-order correlation coefficient *r_xy_*, computed with the scipy.stats.spearmanr function (Virtanen et al., 2020).

## Results

The occurrence of fire promoted biodiversity in both Mediterranean and Boreal communities, as shown for all the four indicators (fig. 1). The median of the distribution of the Species Richness differences Δ*S* (panel A in fig. 1), which excludes null values, resulted significantly greater than zero from the Sign test (whose statistics and p-values are reported in table C1 in Supporting information). Likewise, results from the Wilcoxon signed-rank test (whose statistics and p-values are reported in table C2 in Supporting information), showed that the distributions underlying the biodiversity indicator differences Δ*I*, Δ*F_R_*, and Δ*F_D_* were centered around a value significantly greater than 0 for both ecosystem types (panels B, C, and D in fig. 1). Furthermore, the distributions of Δ*I* resulted skewed towards positive values, meaning that there were more communities with increased levels of biodiversity due to the presence of fire than with reduced levels. These differences are driven by a large diversity increase at intermediate fire return time (see figures in Supporting information). Conversely, at high fire return times (rare fires) the communities have almost the same number of species as they would have without fires, while at short fire return times (frequent fires) the communities are generally less diverse than they would be without fires. These patterns are especially evident for species richness and the inverse Simpson index, while for the functional diversity indices this is more prominent for the Boreal than for the Mediterranean communities (fig. C1).

**Figure 1:**
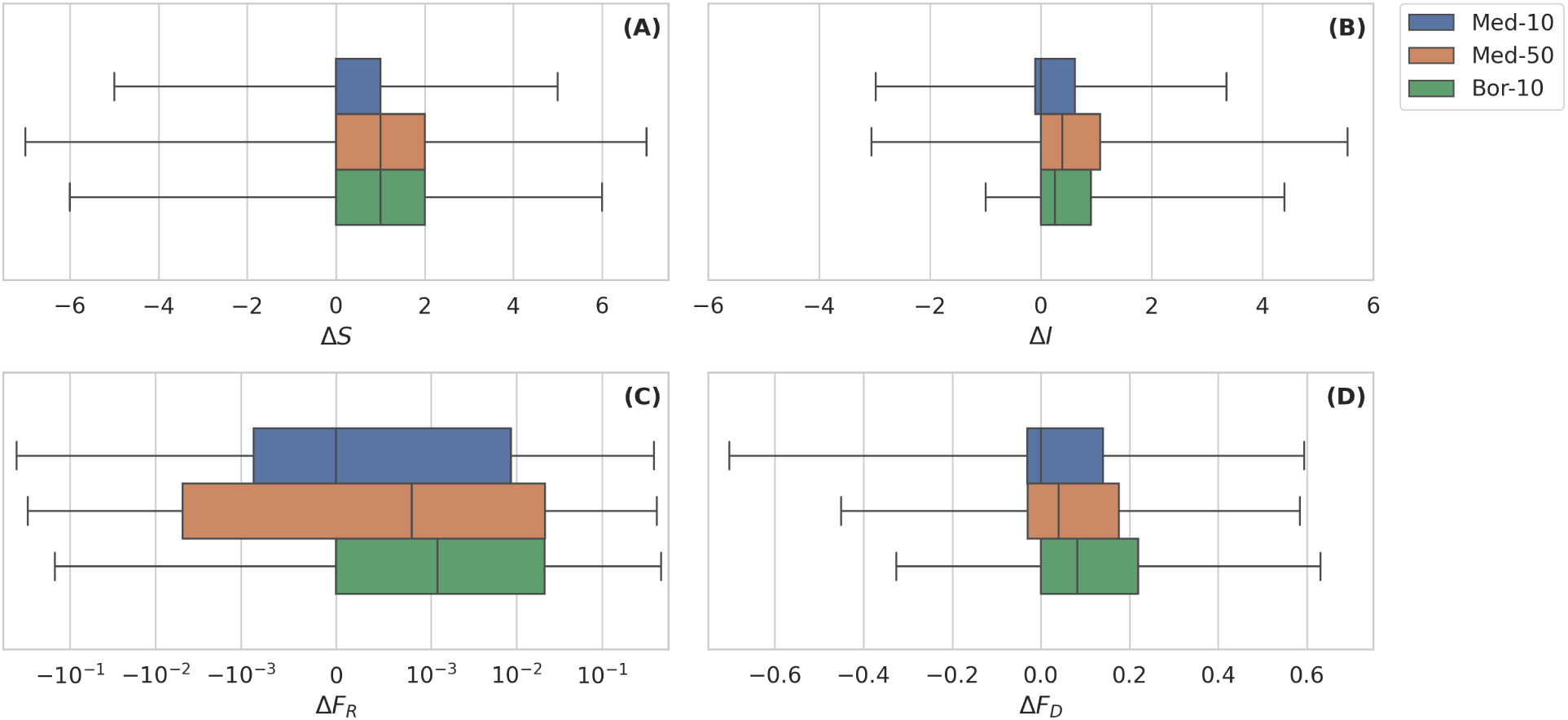
Distributions of the difference between biodiversity indicators for communities with fires (simulated with the fire-vegetation model) and without fires (calculated as equilibria of the Tilman model): A) Species richness (Δ*S*); B) Inverse Simpson Index (Δ*I*); C) Functional Richness (Δ*F_R_*) and D) Functional Divergence (Δ*F_D_*). The colors identify the distributions for the ecosystem types and initial species number (see legend). All of the distributions resulted centered around a value significantly greater than 0 from the Sign or the Wilcoxon signed-rank.

To answer the second research question, plotting the indicators of compositional and functional diversity as a function of the average fire return time, we observed that, although the communities show a wide spread of diversity values, the maximum level of biodiversity occurred at intermediate fire frequency (figs. 2-3). Both compositional diversity indices *S* and *I* reached their maximum value at intermediate return times, and local minimums at the two extremes of the fire axis (fig. 2). We observed this pattern in all three types of simulated communities. Although the polynomial order of the best-fit curves was not identical between quantiles and community types, all curves reflected a hump-shaped pattern. Similar results were obtained when including annual species in the Mediterranean simulations, which also reached larger values of both *S* and *I* at intermediate fire frequencies (Supporting information). Incidentally, we notice that the highest level of Species Richness *S* (fig. 2, top panels) was observed among Med-50 communities (median (med.) of 3 and maximum (max.) of 9 coexisting species; 6 and 16 when including annuals, see fig. H2 in Supporting information), followed by the Bor-10 communities (med. 2, max. 8 coexisting species), and the Med-10 communities (med. 2, max. 6 coexisting species). Similarly, the Inverse Simpson Index *I* values (fig. 2, bottom panels) were highest for the Med-50 communities (med. 1.9, max. 6.8; reaching 2.4 and 10.2 when including annuals, see fig. H2 in Supporting information), followed by the Bor-10 communities (1.4 and 5.4) and the Med-10 communities (1.4 and 4.3). These differences were due to the different size of the initial species simulated, representing the species pool, which led to a better exploration of the parameter space and therefore to the possibility of finding the right combination of species coexisting for larger *N* (Miller et al., 2024).

**Figure 2:**
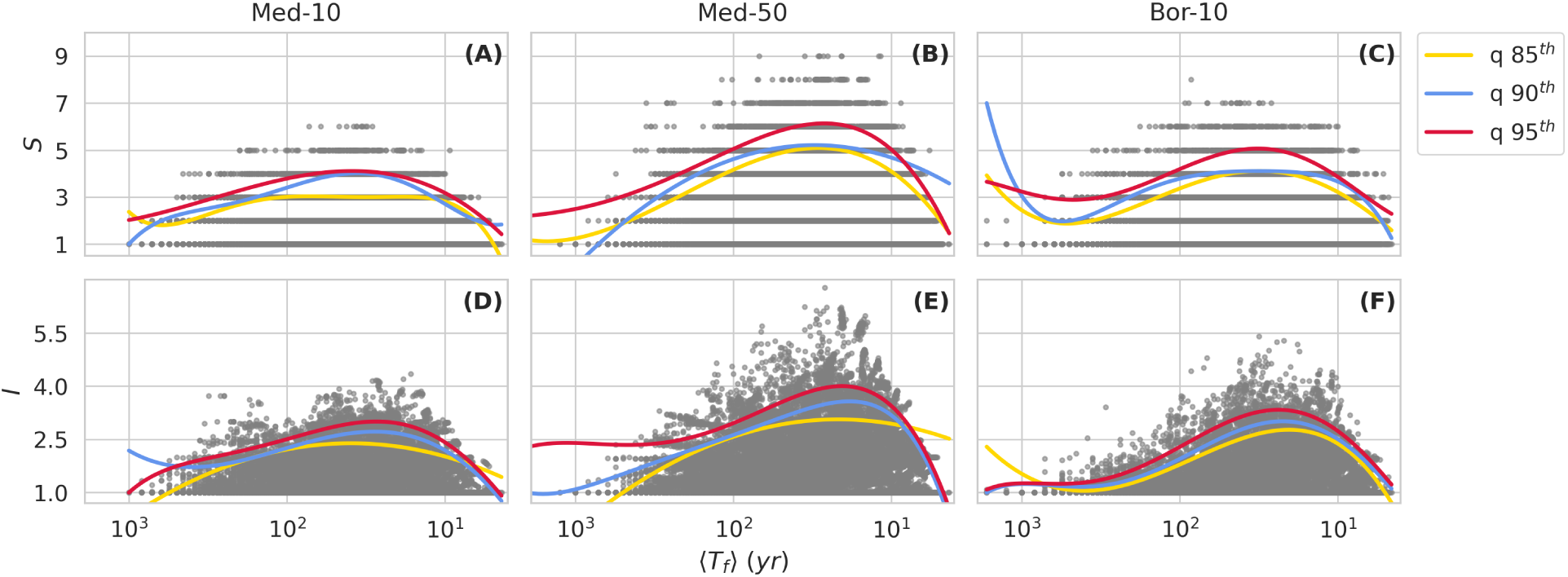
Compositional diversity plots as a function of fire return time for the Mediterranean communities with *N* = 10 (A, D) and *N* = 50 (B, E) initial simulated species, and for the Boreal communities of *N* = 10 (C, F) initially simulated species. Species Richness *S* (A, B, C) and Inverse Simpson Index *I* (D, E, F) are plotted versus average fire return time ⟨*T_f_* ⟩. The 95*^th^* (red), 90*^th^* (blue), and 85*^th^* (yellow) quantile regressions are reported for each plot, and their polynomial order is chosen based on the lowest AIC value (see Tables in Supporting information. The x-axes are reversed to show the increment of the fire frequency, thus going from rare to frequent fires, in logarithmic scale.

**Figure 3:**
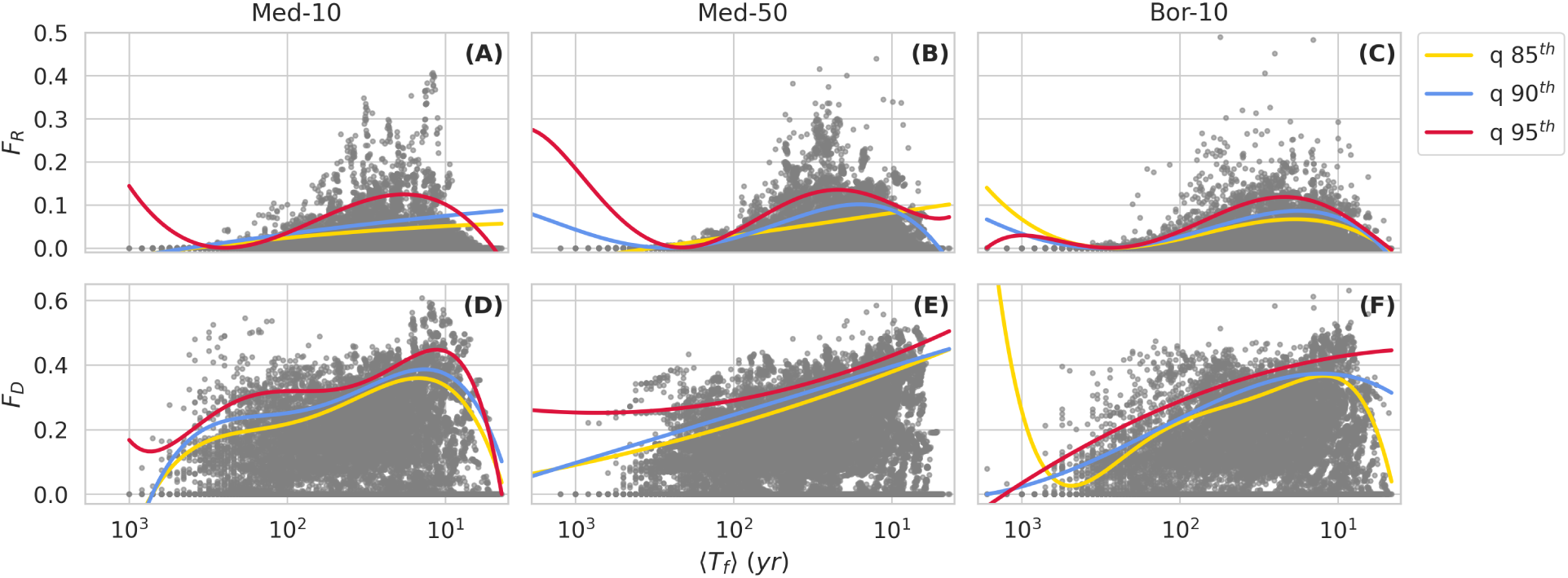
Functional diversity plots over fire frequency for the Mediterranean communities with *N* = 10 (A, D), *N* = 50 (B, E), and for the Boreal communities with *N* = 10 (C, F) initial species. Functional Richness *F_R_* (A, B, C) and Functional Divergence *F_D_* (D, E, F) based on the average plant cover are plotted versus average fire return time ⟨*T_f_* ⟩. The 95*^th^* (red), 90*^th^* (blue), and 85*^th^* (yellow) quantile regressions are reported for each plot, and their polynomial order is chosen based on the lowest AIC value (see tables D7-D12 in Supporting information). Note that the x-axes are reversed to show the increment of the fire frequency as in fig. 2.

Functional Richness *F_R_* (fig. 3, top panels) also generally exhibited maxima occurring at intermediate-high and high fire frequency for all ecosystems. However, some of the lower quan-tile curves did not show the characteristic hump shape: the 85*^th^* quantile regression of the Med-10 and Med-50 communities, as well as the 90*^th^* quantile regression of the Med-10 communities, de-creased linearly towards lower fire frequency regimes. Functional Divergence *F_D_*(fig. 3, bottom panels) showed a broader set of shapes across biomes and experiments, with a predominantly decreasing trend towards low fire frequencies across all cases. Specifically, the quantile regres-sion curves for the Med-10 community types peaked at high frequencies and decreased steeply towards very high frequencies and more slowly towards low frequencies. A similar pattern was observed for the Bor-10 85*^th^* and 90*^th^* quantile regression curves, while the 95*^th^* curve decreased monotonically towards lower fire return time. All the quantile curves of the Med-50 communities also displayed this decreasing trend with fire return time. Similar patterns were found for the Med-50 communities simulated including annuals (see fig. H3 in Supporting information). In all cases, functional diversity reached similar maximum values for both biomes.

As for compositional diversity, the highest levels of functional diversity were observed on average in the Med-50 communities (median of *F_R_*= 0.009 and *F_D_* = 0.18; 0.0 and 0.2 when including annuals, see Supporting information), followed by the Bor-10 (*F_R_* = 0.003 and *F_D_* = 0.13) and the Med-10 communities (*F_R_* = 0.001 and *F_D_* = 0.12). Conversely, Bor-10 presented the maximum value of Functional Richness (0.49) and Divergence (0.77). Nevertheless, for both functional diversity indicators, the maximum values of the distributions resulted more similar across ecosystems (relative difference of 16% for *F_R_* and 8% for *F_D_*) as compared to the compositional diversity indicator (relative difference of 33% for *S* and 38% for *I*).

Comparing fig. 3 with fig. 2, we note that the quantile regression curves corresponding to functional diversity exhibited more dissimilarity between community types and metrics as compared to the compositional diversity curves. We also note that functional richness exhibited an elevated number of communities with low diversity values compared to the other biodiversity indicators. This was mainly due to the geometric dependency of *F_R_* on *S*, as in the 3-dimensional parameter space, communities with species richness *S* ≤ 3 created hyperplanes rather than hypervolumes, whose volume was equal to or close to 0. Finally, it is worth mentioning that the sharp and increasing trend of some of the fit lines at low fire frequency (high fire return time, fig. 2 C, fig. 3 A, B, C, and F) is not relevant, given it was caused by a few isolated points which heavily influenced the shape of the curve fit.

To answer the third research question, we analyzed whether a larger number of coexisting plant species was associated with a larger variety of functional strategies (as captured by our model parameters). All biodiversity indicators positively and significantly correlated with each other in the different community types (see Supporting information), with the strongest correlations found between indicators of the same kind (compositional and functional). The correlations of Species Richness with Functional Richness resulted strong and with similar magnitudes as other indices combinations (*r* ≥ 0.6), while the relationships with Functional Divergence were weaker, with *r* ≤ 0.4 in all the community types. Indeed, the maxima of functional diversity were not observed at the maximum values of specific richness (fig. 4), showing that the communities with the largest number of species were not those with the greatest functional diversity. This suggests that, in the presence of fire, some level of similarity between species allowed for a greater number of coexisting species.

**Figure 4:**
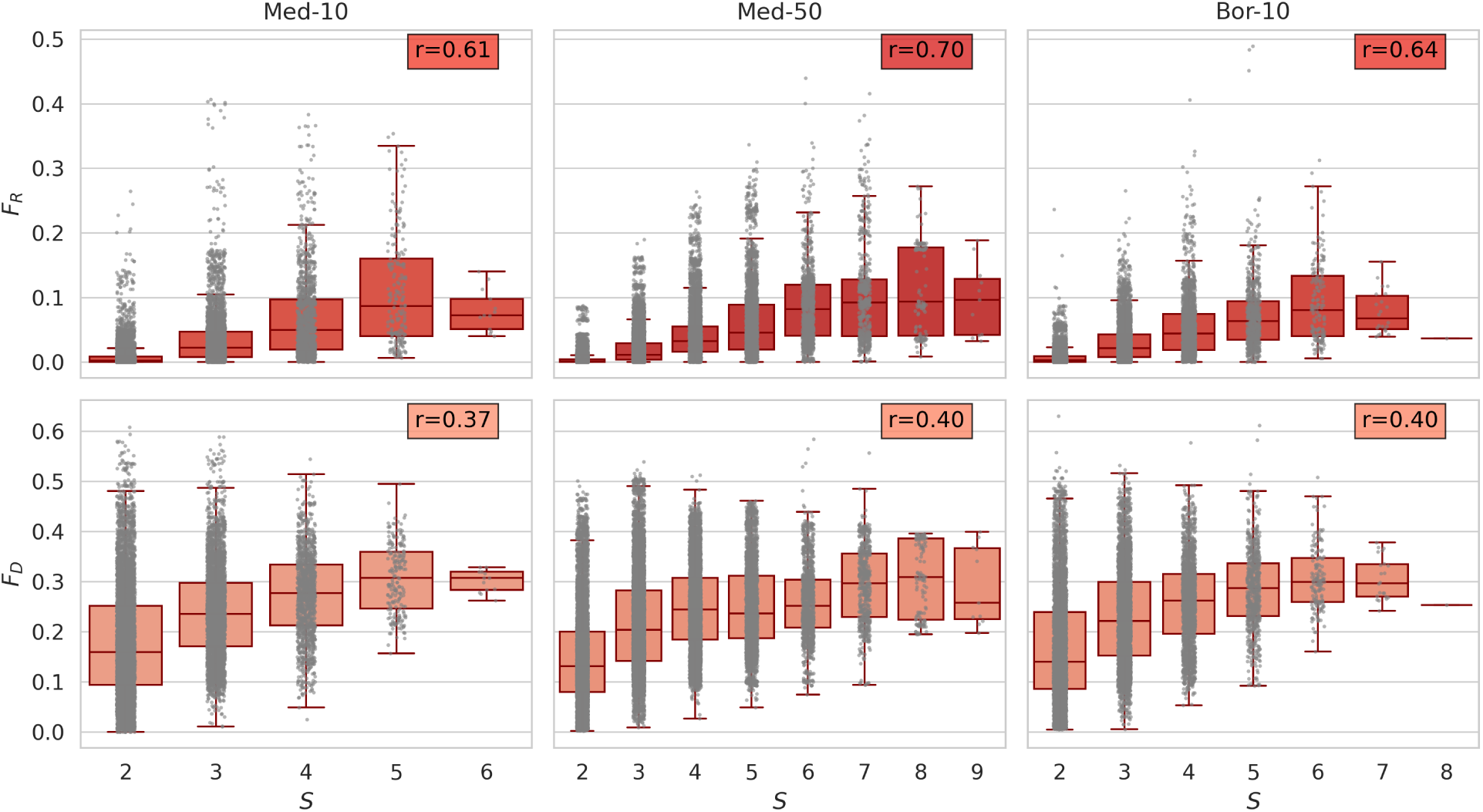
Functional Richness (top panels) and Functional Divergence (bottom panels) boxplot with jitter for different values of Species Richness, for: Mediterranean *N* = 10 (left panels); Mediterranean *N* = 50 (central panels); Boreal *N* = 10 (right panels). The Spearman rank-order correlation coefficients *r* are reported for each panel and they all resulted in significant correlation (*p <* 0.001). Nonetheless, the scatter plots show that the community with the highest number of species does not coincide with those with the largest functional diversity indices.

## Discussion

Our model results showed that, for both Mediterranean and Boreal fire-prone ecosystems, plant diversity peaked at medium fire frequencies and decreased towards the extreme of the fire return time gradient. While the biodiversity indicators analyzed (Species Richness, Inverse Simpson Index, Functional Richness, and Divergence) were quite scattered when plotted as a function of fire return time, their top quantiles followed a hump-shaped curve as a function of fire return time in most cases. Compositional and functional diversity were generally higher in the simulations of plant communities prone to fires, as compared to simulations of the same set of species without fires. Furthermore, the highest plant community diversity occurred at intermediate fire frequencies, while less diverse communities were observed towards the extremes of the fire frequency gradient. Yet, the two types of diversity did not peak in the same communities.

On average, fires tended to increase diversity, as observed by comparing the simulated communities obtained with and without fires. Furthermore, the highest levels of diversity and the largest gain in diversity were at intermediate values of fire frequency, but closer to the frequent end. This is consistent with the high biodiversity reported for fire-prone systems (He et al., 2019;

Pausas and Ribeiro, 2017) and their tendency to maintain greater diversity under frequent burn-ing conditions than under fire exclusion conditions (e.g. Buness et al., 2025; Fernandez-Garcia et al., 2020; Pellegrini et al., 2023). We found the lowest biodiversity levels in low-frequency fire systems, where single species survived alone, similarly to real ecosystems dominated by late-successional species (e.g. Amici et al., 2013). Remarkably, our study showed that functional diversity generally peaked at intermediate fire frequencies, while these patterns are most often shown only for compositional diversity.

Our results thus followed the predictions of the Intermediate Disturbance Hypothesis: at infrequent fires, communities were likely to be late-successional, competition-dominated communities, as we showed that the simulated diversity of communities with an identical species pool with and without fires was very similar. Conversely, at high fire frequency, the fires reduce diversity as only a handful of species can survive in such high frequency conditions. In between these two extremes, an optimum is observed, where we observed the largest gain in diversity. Mixed evidence has been reported for the IDH in general, as it fails to describe numerous empirical and theoretical results (Mackey and Currie, 2001), and has been criticized theoretically for a lack of appropriate mechanistic explanation behind the expected pattern (Fox, 2013). The IDH has been observed for fire-prone plant communities along a fire frequency gradient, although not in all instances (e.g. He et al., 2019; Pellegrini et al., 2023). The emergence of the IDH in fire-prone plant communities might be related to the vegetation-fire feedback, which is included in our model and is very relevant in real communities (e.g. Baudena et al., 2015; Pausas and Dantas, 2017; Rohde, 2013). In fact, in a recent paper with a model rather similar to ours (Liao et al., 2022, based on the same competition-colonization equations and including fires as pulse disturbances), the IDH was not observed for plant compositional diversity. This difference possibly arises because in Liao et al. (2022) all the species were assumed to be affected to the same extent by fires — that is, the fire response was set equal for all species — or because fires were represented as deterministic events, without including any vegetation-fire feedback. This latter explanation seems to be supported by an experiment we performed with a simplified version of our model, where we excluded the fire-vegetation feedback (by including stochastic fires with a fixed average return time). The simulations did not display a maximum nor a hump shape for diversity indices, but, similarly to Liao et al. (2022), a multimodal relationship emerged when plotting data for specific communities along an imposed fire frequency gradient (see fig. G1 in Supporting information). This result is also consistent with the seminal work of Zinck and Grimm (2009), which showed that models including the plant-fire feedback (which they named “regrowth-dependent flammability”) could produce the IDH patterns, although using simpler diversity indicators – namely, the number of successional stages in a landscape. Given the im- portance of the fire-vegetation feedback for different ecosystems (Pausas and Dantas, 2017), due to the effects of vegetation flammability on fire spread and the importance of the different post-fire regrowth strategies (Archibald et al., 2018; Harrison et al., 2021; Pausas and Verdú, 2008; Simpson et al., 2022) our results are likely to represent a key feature of many fire ecosystems.

The compositional and functional diversity indices were correlated with one another, with stronger correlation between species richness and functional richness than with functional divergence, similar to what was observed in biogeographical studies (Ordonez and Svenning, 2015). This is partly expected from the functional diversity indices used, although the correlation is less likely in modeled ecosystems (such as ours, see Magnani et al., 2023) where differences in assembly order can lead to different final community composition (Mason et al., 2013). Yet, the two types of diversity were also partly decoupled, similarly to observational studies in different biomes (e.g. Mirabel et al., 2020; Navarro-Cano et al., 2024; Pellegrini et al., 2023; Swenson et al., 2012): high levels of compositional and functional biodiversity were not observed in the same plant communities, although for both metrics the peaks occurred at similar fire frequencies. This suggests that species that coexist in fire-prone ecosystems are not necessarily very different and might in fact share some characteristics, as fire selects these traits, being a strong community assembly process, as previously suggested by Pausas and Verdú (2008): for example, species might have a comparable growth rate and similar performance in fire response, enabling them to survive the same fire regime. Finally, it is worth mentioning that, since our model does not represent evolution nor its timescales, we did not investigate phylogenetic diversity, which is a fundamental component of diversity (Molina-Venegas et al., 2022).

Our approach allowed us to make a general statement about the fire-diversity relationship by examining different parameterized ecosystems and considering different possible species pools associated with their self-sustained fire regime. We simulated the Mediterranean with a much larger species pool than the Boreal ecosystems. However, species richness of the Mediterranean communities was smaller than what was expected from the observed plant diversity in the Mediterranean Basin biome (Cowling et al., 1996; Myers et al., 2000), although including annuals solved this issue (fig. H2 in Supporting information). The relatively low values of species richness in the Mediterranean ecosystem might also be due to an undersampling of the simulated communities, since, despite the large number of simulations performed, only a portion of the possible parameter combinations was explored, given the large dimensionality of the parameter space, especially for the simulations with 50 initial species. We also note that functional diversity was similar in the Mediterranean and Boreal biomes, despite the much higher species richness in the former (especially when annuals are included; see Supporting information). This pattern may reflect the stronger functional clustering of species in the Mediterranean (Molina-Venegas et al., 2022).

Future development could disentangle which plant characteristic let the hump-shaped distribution emerge in the model dynamics, showing, for example, which types of plant functional responses or characteristics of the initial species set (representing the species pool) are necessary for the final communities to be more or less diverse. Analyzing the parameter variability would help to understand the increase in functional diversity when including the fire-feedback, especially for the Boreal ecosystems. These studies could build on our current understanding that the fire-vegetation feedback, due to fire-related plant traits that determine plant flammability and the ability to re-grow after fires, is determinant not only for ecosystem resilience (Baudena et al., 2020; Magnani et al., 2023) but also for ecosystem compositional and functional diversity.

The connection between fires and biodiversity is also very relevant from a management perspective, for example, in relation to fire suppression or prescribed-burning strategies (e.g. Puig-Gironès et al., 2025). More generally, to predict the fate of ecosystems subject to global change, affecting biodiversity and wildfire regimes, and to support restoration action and management, modeling efforts should crucially include the vegetation-fire feedback, which, however, is often only partly represented in most global vegetation models (Hantson et al., 2016; Harrison et al., 2021; Kelley et al., 2014; Venevsky et al., 2019).

## Acknowledgments

We acknowledge financial support under the National Recovery and Resilience Plan (NRRP), Mission 4, Component 2, by the Italian Ministry of University and Research (MUR), funded by the European Union - NextGenerationEU, specifically: i) M.T. was supported by Investment 3.3 (D.M. 352, 09/04/2022). ii) G.V. and M.M. were supported by the Italian National Biodiversity Future Center (NBFC), Investment 1.4 (Project code CN00000033). iii) M.B. was supported by Investment 1.1 Call for tender No. 1409 published on 14.9.2022, with the PRIN-PNRR 2022 project Plant traits of native and invasive species in fire ecosystems across the world (WiFIN; P2022NRLF2; CUP B53D23023670001). R.D.S. was supported by the Spanish MICINN (PID2022-138158OB-I00).

The authors are grateful to Gianluigi Ottaviani for references and explanations about functional diversity patterns, and to Sara Bernardi for useful suggestions on the mathematical approach.

## Statement of Authorship

R.D.S. and M.B. conceived the original idea; M.T. and G.V. developed the software (in Python and Fortran, respectively); G.V. led the model parametrization and simulations; M.T. led the analyses and visualization of results; R.D.S., M.M. and M.E. provided useful feedback during the analyses; M.T., G.V. and M.B. wrote the original draft; M.B. supervised the work; All the authors contributed to reviewing and editing.

## Data and Code Availability

The data shown in the manuscript were produced by model simulations. The simulation code and analysis methods for the reproduction of results presented are uploaded as zip file, and will be available as a GitHub repository upon acceptance. available in the GitHub repository: https://github.com/torrassam/Fire-Diversity.git.

