## Supplementary Information for "Functional and compositional diversity peak at intermediate fire frequencies when modeling the plant-fire feedback"

#### Table of Contents:

**A Model Parameters** **2**

**B Hypervolume visualization** **4**

**C Comparing biodiversity with and without fires** **5**

**D Quantile Regression** **9**

D.1 Compositional Diversity . . . . . . . . . . . . . . . . . . . . . . . . . . . . . . . . 9

D.2 Functional Diversity . . . . . . . . . . . . . . . . . . . . . . . . . . . . . . . . . . 15

**E Biodiversity indices distributions** **21**

**F Biodiversity indices correlations** **22**

**G Simulations with fixed average fire return time** **23**

**H Mediterranean simulations including annual grasses** **24**

H.1 Extending the range of $c$ values to annual species . . . . . . . . . . . . . . . . . . 24

H.2 Compositional and Functional diversity with and without annual species . . . . . . 25

### A Model Parameters

Table A1: Reference parameters of colonization and mortality for Mediterranean communities. Values from Baudena et al. (2020)

| **Hierarchy** | **Colonization rate** [$y^{-1}$] | **Mortality rate** [$y^{-1}$] |
|---|---|---|
| 1 | 0.047 | 1/400 |
| 2 | 0.053 | 1/125 |
| 3 | 0.045 | 1/50 |
| 4 | 0.067 | 1/25 |
| 5 | 0.11 | 1/15 |
| 6 | 0.22 | 1/40 |

Table A2: Reference parameters of colonization and mortality for Boreal communities. Values from Magnani et al. (2023)

| **Hierarchy** | **Colonization rate** [$y^{-1}$] | **Mortality rate** [$y^{-1}$] |
|---|---|---|
| 1 | 0.085 | 0.035 |
| 2 | 0.13 | 0.015 |
| 3 | 0.17 | 0.023 |

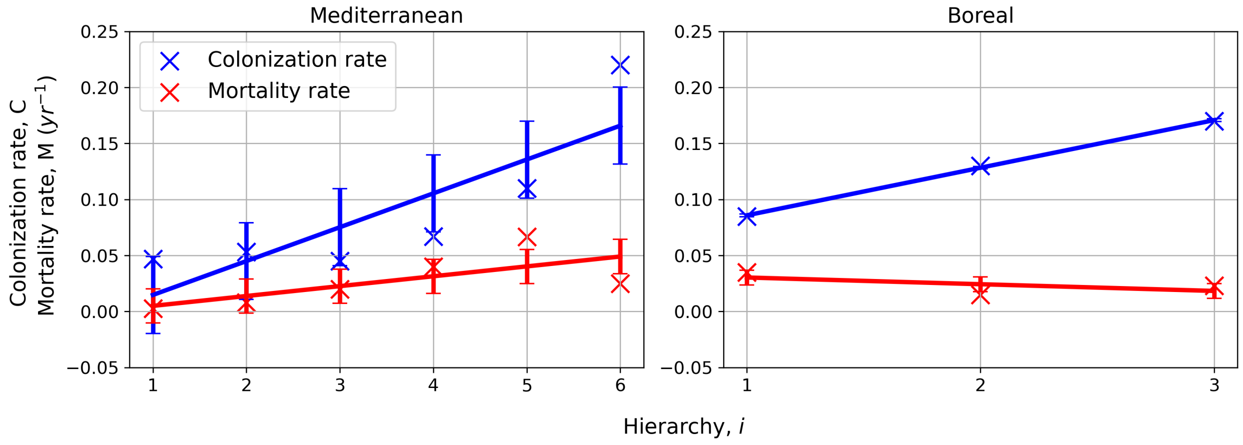


Figure A1: Linear regressions of colonization $C$ and mortality rates $M$ with respect to plant position in the competition hierarchy $i$. Left panel: Mediterranean ecosystem. Right panel: Boreal ecosystem.

Table A3: Parameters found with regressions.

| **Parameters [$y^{-1}$]** | **Mediterranean ecosystem** | **Boreal ecosystem** |
|---|---|---|
| $\alpha_C$ | 0.03023 | 0.0425 |
| $\beta_C$ | -0.01547 | 0.0433 |
| std($\gamma_{C_i}$) | 0.03435 | 0.0012 |
| $\alpha_M$ | 0.00881 | -0.0060 |
| $\beta_M$ | -0.00382 | 0.0363 |
| std($\gamma_{M_i}$) | 0.01526 | 0.0066 |

### B Hypervolume visualization

In fig. B1 , we show the representation of a hypervolume in the trait space, retrieved from a Med-50 community composed as reported in Table B1.

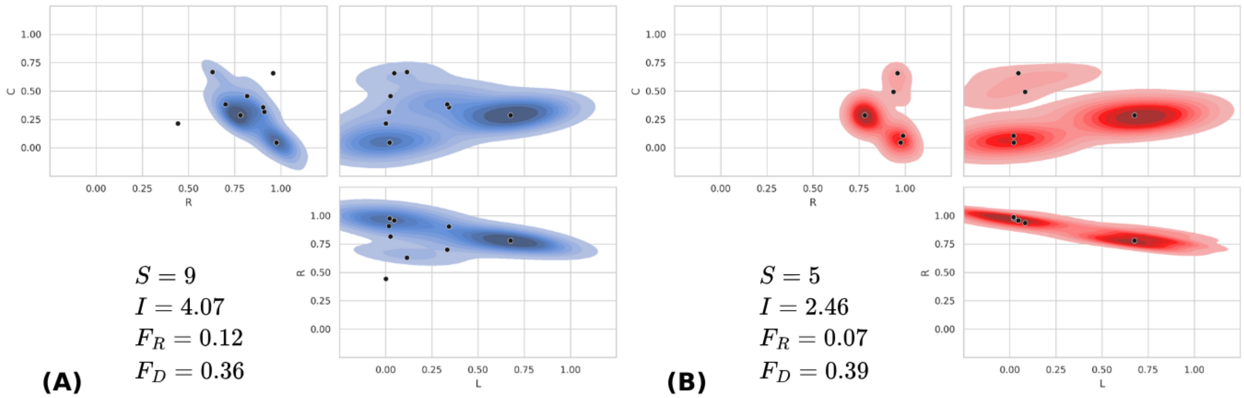


Figure B1: Hypervolume representation of two resulting Med-50 communities in the $C$-$R$-$L$ trait space. Black dots are the positions in the trait space of the surviving species.

Table B1: Species traits and composition of the Med-50 communities (A and B) whose hypervolumes are displayed in fig. B1.

| $i$ | $C_i$ | $M_i$ | $R_i$ | $L_i$ | $\langle b_i \rangle^{(A)}$ | $\langle b_i \rangle^{(B)}$ |
|---|---|---|---|---|---|---|
| 1 | 0.013650 | 0.009410 | 0.976090 | 0.012580 | 0.174 | 0.141 |
| 4 | 0.064120 | 0.011290 | 0.442830 | 0.002680 | 0.007 | - |
| 5 | 0.031530 | 0.022310 | 0.987030 | 0.011740 | - | 0.050 |
| 15 | 0.086570 | 0.012910 | 0.780940 | 0.338220 | 0.231 | 0.344 |
| 19 | 0.095200 | 0.017340 | 0.910120 | 0.010290 | 0.041 | - |
| 22 | 0.107390 | 0.018000 | 0.903760 | 0.172430 | 0.056 | - |
| 26 | 0.136910 | 0.025330 | 0.815680 | 0.014680 | 0.027 | - |
| 27 | 0.115050 | 0.007780 | 0.699570 | 0.167820 | 0.060 | - |
| 36 | 0.197190 | 0.038460 | 0.957300 | 0.025030 | - | 0.035 |
| 37 | 0.197190 | 0.038460 | 0.957300 | 0.025030 | 0.004 | 0.022 |
| 40 | 0.200670 | 0.015220 | 0.630040 | 0.059280 | 0.016 | - |

Note: For each surviving species, its traits and average vegetation covers is reported.

### C Comparing biodiversity with and without fires

We here study the distributions of the differences between biodiversity indicators estimated for the same communities (i.e., same set of initial parameters) with and without fires, the latter being the standard competition-colonization Tilman model (1994). To evaluate whether their means were significantly different from zero, we performed statistical tests separately for each biome and its subsets (Med-10, Med-50, and Bor-10). Each of these distributions comprised a total number of points equal to the number of simulations in the fire-vegetation feedback model (100.000). The following two tables report the results of these tests. These are followed by two figures representing the four indicators as a function of fire return time, with and without fires, for all the different biomes and experiments.

Table C1: Results of the Signed test for the distribution of the differences of Species Richness $\Delta S$.

| **Biodiversity indicator** | **Ecosystem type** | **M** | **$p$-value** |
|---|---|---|---|
| | Med-50 | 4976 | 0.0 |
| $\Delta S$ | Med-10 | 1352 | 0.0 |
| | Bor-10 | 3918 | 0.0 |

Note: The Alternative Hypothesis tested is that the median of the distribution of the differences is greater than zero ($\Delta S > 0$). The $M$-statistic is estimated as $M = (N(+) - N(-))/2$, where $N(+)$ and $N(-)$ are respectively the number of values lower and greater than 0.

Table C2: Results of the Wilcoxon signed-rank test for the distribution of the differences of biodiversity indicators: Inverse Simpson Index $\Delta I$, Functional Richness $\Delta F_R$, and Functional Divergence $\Delta F_D$)

| **Biodiversity indicator** | **Ecosystem type** | **U-statistic** | ***p*-value** |
|---|---|---|---|
| $\Delta I$ | Med-50 | 361719713 | 0.0 |
| | Med-10 | 65998867 | 0.0 |
| | Bor-10 | 78120705 | 0.0 |
| $\Delta F_R$ | Med-50 | 280410330 | 0.0 |
| | Med-10 | 60355548 | 0.0 |
| | Bor-10 | 70301374 | 0.0 |
| $\Delta F_D$ | Med-50 | 330491312 | 0.0 |
| | Med-10 | 64957723 | 0.0 |
| | Bor-10 | 76840023 | 0.0 |

Note: The Alternative Hypothesis tested is that the distribution of such differences is stochastically greater than a distribution symmetric about zero ($\Delta > 0$). The $U$-statistic is estimated as the sum of the ranks of the differences greater than 0.

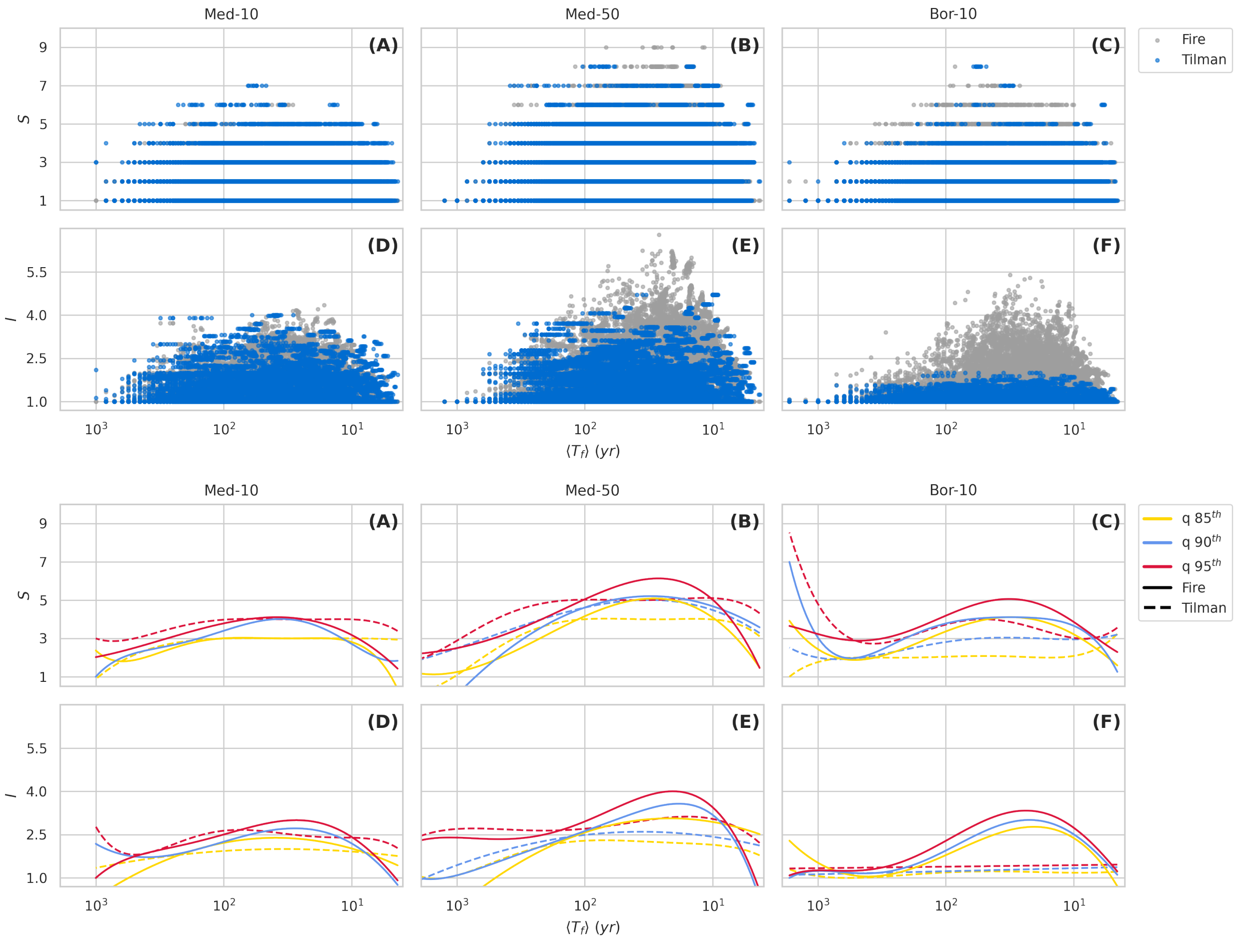


Figure C1: Compositional diversity plots as a function of fire return time for all ecosystem types modeled with and without fires (Tilman model). Top panels: scatterplots for the model with fires (gray) and without fires (blue). Bottom panels: $95^{th}$ (red), $90^{th}$ (blue), and $85^{th}$ (yellow) quantile regression lines, with polynomial order based on the lowest AIC value, for the model with fires (solid lines) and without fires (dashed lines).

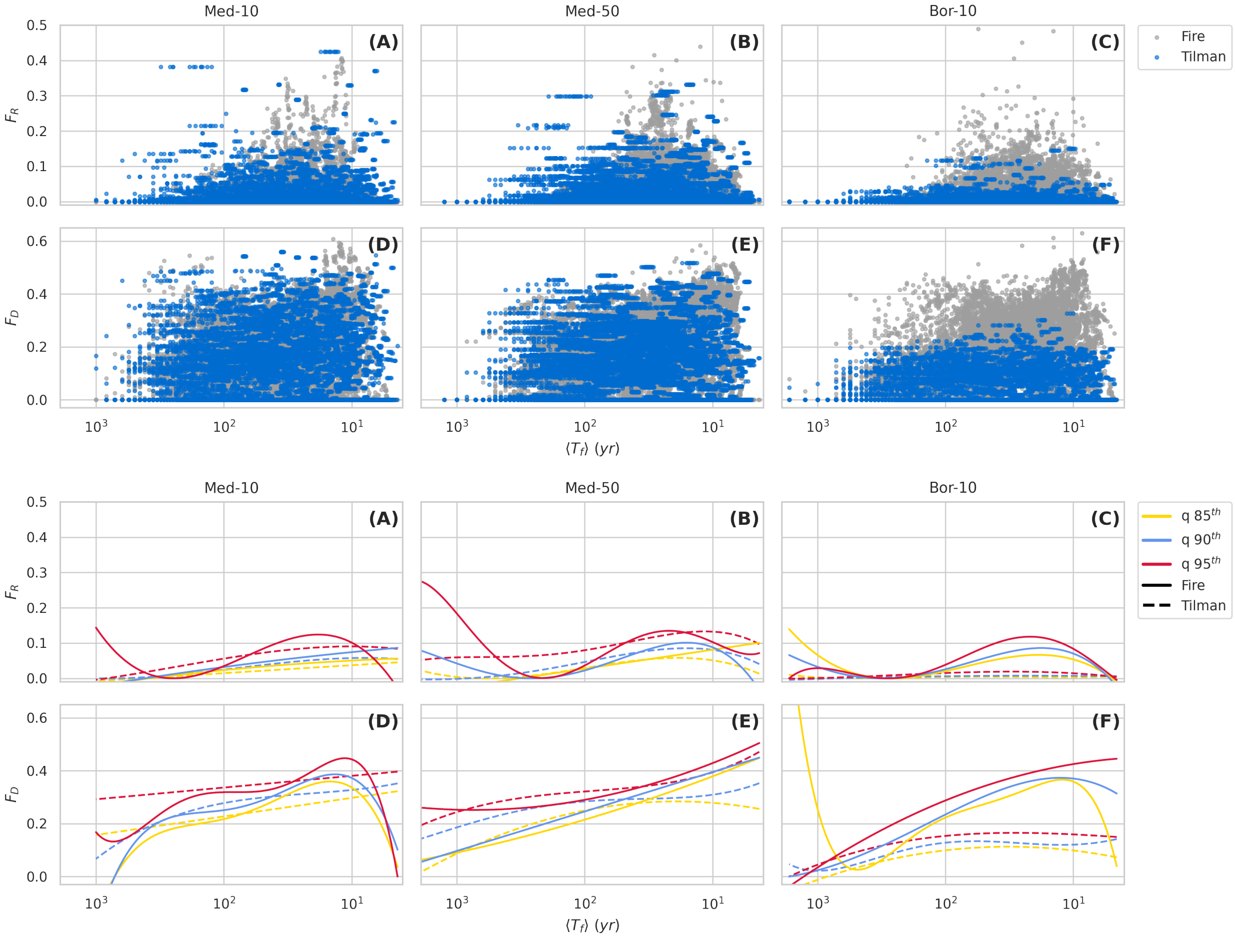


Figure C2: Functional diversity plots as a function of fire return time for all ecosystem types modeled with and without fires (Tilman model). Top panels: scatterplots for the model with fires (gray) and without fires (blue). Bottom panels: $95^{th}$ (red), $90^{th}$ (blue), and $85^{th}$ (yellow) quantile regression lines, with polynomial order based on the lowest AIC value, for the model with fires (solid lines) and without fires (dashed lines).

### D Quantile Regression

To select the best polynomial order for $85^{th}$, $90^{th}$, and $95^{th}$ quantile regression for the diversity-disturbance plot, we have computed the Akaike Information Criterion (AIC), pseudo R-squared, Wald test statistic, and p-value. The Wald test is used to assess the significance of an individual or group of coefficients in regression models Koenker and Portnoy (1996): the null hypothesis that all coefficients resulting from the polynomial regression fit are equal to zero is tested. By obtaining a test p-value $< 0.05$, we can reject the null hypothesis with 95% confidence and accept the alternative hypothesis that not all predictor variables in the set of coefficients are equal to zero.

#### *D.1 Compositional Diversity*

Table D1: Quantile regression statistics of the Species Richness and fire return time relationship for the $N = 10$ Mediterranean simulations (Med-10).

| **Quantile** | **Polynomial Order** | **AIC** | **R-squared** | **Wald test Statistics** | **Wald test p-value** |
|---|---|---|---|---|---|
| 0.85 | 1 | 63938.7 | -0.000 | 0.0 | 1.0 |
| | 2 | 61742.4 | 0.015 | 1368.8 | 0.0 |
| | 3 | 61563.4 | 0.019 | 1253.4 | 0.0 |
| | 4 | 61517.7 | 0.024 | 9466.5 | 0.0 |
| | 5 (*) | 60769.5 | 0.028 | 10008.7 | 0.0 |
| 0.90 | 1 | 63938.7 | -0.000 | 0.0 | 1.0 |
| | 2 | 72587.7 | 0.057 | 2743.6 | 0.0 |
| | 3 | 72265.8 | 0.069 | 3462.6 | 0.0 |
| | 4 | 71220.0 | 0.073 | 4520.0 | 0.0 |
| | 5 (*) | 71146.7 | 0.074 | 3971.5 | 0.0 |
| 0.95 | 1 | 81411.0 | -0.000 | 0.0 | 1.0 |
| | 2 | 76141.0 | 0.059 | 4359.6 | 0.0 |
| | 3 (*) | 75857.2 | 0.069 | 5264.5 | 0.0 |
| | 4 | 75887.5 | 0.070 | 5241.0 | 0.0 |
| | 5 | 75988.9 | 0.070 | 5757.3 | 0.0 |

Note: (*) indicates the best polynomial order for each quantile.

Table D2: Quantile regression statistics of the Inverse Simpson Index and fire return time relationship for the $N = 10$ Mediterranean simulations (Med-10).

| Quantile | Polynomial Order | AIC | R-squared | Wald test Statistics | Wald test p-value |
|---|---|---|---|---|---|
| 0.85 | 1 | 47937.4 | 0.013 | 361.5 | 0.0 |
| | 2 (*) | 46895.4 | 0.064 | 475.0 | 0.0 |
| | 3 | 47130.3 | 0.084 | 1191.2 | 0.0 |
| | 4 | 47123.7 | 0.084 | 1268.3 | 0.0 |
| | 5 | 47338.0 | 0.086 | 1399.1 | 0.0 |
| 0.90 | 1 | 55756.6 | 0.015 | 593.2 | 0.0 |
| | 2 | 53598.6 | 0.070 | 419.2 | 0.0 |
| | 3 | 53529.7 | 0.094 | 809.0 | 0.0 |
| | 4 (*) | 53450.7 | 0.094 | 815.8 | 0.0 |
| | 5 | 53499.0 | 0.099 | 1488.7 | 0.0 |
| 0.95 | 1 | 64330.8 | 0.013 | 533.9 | 0.0 |
| | 2 | 62104.2 | 0.069 | 687.4 | 0.0 |
| | 3 | 60742.4 | 0.092 | 1771.4 | 0.0 |
| | 4 | 60845.8 | 0.092 | 1636.9 | 0.0 |
| | 5 (*) | 60675.4 | 0.093 | 1901.2 | 0.0 |

Note: (*) indicates the best polynomial order for each quantile.

Table D3: Quantile regression statistics of the Species Richness and fire return time relationship for the $N = 50$ Mediterranean simulations (Med-50).

| **Quantile** | **Polynomial Order** | **AIC** | **R-squared** | **Wald test Statistics** | **Wald test p-value** |
|---|---|---|---|---|---|
| 0.85 | 1 | 145082.5 | 0.003 | 787.1 | 0.0 |
| | 2 | 141751.9 | 0.068 | 4758.9 | 0.0 |
| | 3 (*) | 140133.8 | 0.078 | 11056.1 | 0.0 |
| | 4 | 140174.9 | 0.078 | 11118.8 | 0.0 |
| | 5 | 141151.5 | 0.079 | 14486.0 | 0.0 |
| 0.90 | 1 | 153043.0 | -0.000 | 0.0 | 1.0 |
| | 2 (*) | 147841.9 | 0.033 | 1502.2 | 0.0 |
| | 3 | 148219.5 | 0.039 | 1184.3 | 0.0 |
| | 4 | 148768.4 | 0.039 | 1048.9 | 0.0 |
| | 5 | 148319.8 | 0.045 | 114125.8 | 0.0 |
| 0.95 | 1 | 173450.3 | -0.000 | 0.0 | 1.0 |
| | 2 | 163851.3 | 0.063 | 2289.6 | 0.0 |
| | 3 | 162627.5 | 0.080 | 5309.1 | 0.0 |
| | 4 (*) | 162559.4 | 0.081 | 5464.9 | 0.0 |
| | 5 | 162842.2 | 0.081 | 6033.0 | 0.0 |

Note: (*) indicates the best polynomial order for each quantile.

Table D4: Quantile regression statistics of the Inverse Simpson Index and fire return time relationship for the $N = 50$ Mediterranean simulations (Med-50).

| **Quantile** | **Polynomial Order** | **AIC** | **R-squared** | **Wald test Statistics** | **Wald test p-value** |
|---|---|---|---|---|---|
| 0.85 | 1 | 106714.8 | 0.030 | 1681.9 | 0.0 |
| | 2 (*) | 103912.1 | 0.055 | 1089.8 | 0.0 |
| | 3 | 104730.9 | 0.074 | 4006.3 | 0.0 |
| | 4 | 104556.5 | 0.076 | 3609.7 | 0.0 |
| | 5 | 104309.0 | 0.078 | 2642.0 | 0.0 |
| 0.90 | 1 | 116943.9 | 0.036 | 2173.9 | 0.0 |
| | 2 | 115655.5 | 0.056 | 836.8 | 0.0 |
| | 3 | 115690.6 | 0.082 | 3128.0 | 0.0 |
| | 4 | 115357.2 | 0.085 | 2690.3 | 0.0 |
| | 5 (*) | 115033.3 | 0.086 | 2342.8 | 0.0 |
| 0.95 | 1 | 132785.8 | 0.034 | 1458.3 | 0.0 |
| | 2 | 131053.5 | 0.056 | 472.9 | 0.0 |
| | 3 | 129815.7 | 0.080 | 1489.9 | 0.0 |
| | 4 (*) | 129694.3 | 0.084 | 1474.7 | 0.0 |
| | 5 | 129950.2 | 0.085 | 1226.5 | 0.0 |

Note: (*) indicates the best polynomial order for each quantile.

Table D5: Quantile regression statistics of the Species Richness and fire return time relationship for the $N = 10$ Boreal simulations (Bor-10).

| **Quantile** | **Polynomial Order** | **AIC** | **R-squared** | **Wald test Statistics** | **Wald test p-value** |
|---|---|---|---|---|---|
| 0.85 | 1 | 70767.3 | 0.022 | 1207.1 | 0.0 |
| | 2 | 68179.1 | 0.101 | 3083.6 | 0.0 |
| | 3 | 66665.1 | 0.118 | 5622.4 | 0.0 |
| | 4 (*) | 66635.2 | 0.119 | 5835.0 | 0.0 |
| | 5 | 66741.3 | 0.120 | 6323.0 | 0.0 |
| 0.90 | 1 | 76140.3 | -0.000 | 0.0 | 1.0 |
| | 2 | 71289.4 | 0.057 | 2474.1 | 0.0 |
| | 3 | 70658.7 | 0.069 | 1915.2 | 0.0 |
| | 4 | 70682.0 | 0.069 | 1866.7 | 0.0 |
| | 5 (*) | 70570.1 | 0.070 | 7896.9 | 0.0 |
| 0.95 | 1 | 83089.8 | 0.022 | 420.7 | 0.0 |
| | 2 | 81313.2 | 0.093 | 942.8 | 0.0 |
| | 3 | 79980.1 | 0.115 | 2559.3 | 0.0 |
| | 4 | 79696.2 | 0.116 | 3157.4 | 0.0 |
| | 5 (*) | 79336.9 | 0.117 | 3616.3 | 0.0 |

Note: (*) indicates the best polynomial order for each quantile.

Table D6: Quantile regression statistics of the Inverse Simpson Index and fire return time relationship for the $N = 10$ Boreal simulations (Bor-10).

| Quantile | Polynomial Order | AIC | R-squared | Wald test Statistics | p-value |
|---|---|---|---|---|---|
| 0.85 | 1 | 48820.6 | 0.104 | 3826.2 | 0.0 |
| | 2 | 47136.9 | 0.124 | 1161.2 | 0.0 |
| | 3 | 46176.6 | 0.202 | 10158.9 | 0.0 |
| | 4 (*) | 46105.6 | 0.202 | 9585.7 | 0.0 |
| | 5 | 46120.9 | 0.205 | 5177.4 | 0.0 |
| 0.90 | 1 | 54716.2 | 0.095 | 3682.4 | 0.0 |
| | 2 | 53709.2 | 0.124 | 1093.9 | 0.0 |
| | 3 | 51590.9 | 0.199 | 4475.3 | 0.0 |
| | 4 | 51571.3 | 0.199 | 4505.5 | 0.0 |
| | 5 (*) | 51544.3 | 0.201 | 4793.1 | 0.0 |
| 0.95 | 1 | 63477.8 | 0.072 | 2937.8 | 0.0 |
| | 2 | 60422.0 | 0.108 | 592.2 | 0.0 |
| | 3 | 59094.8 | 0.177 | 2393.3 | 0.0 |
| | 4 | 59144.3 | 0.178 | 2332.7 | 0.0 |
| | 5 | 59085.7 | 0.178 | 2413.7 | 0.0 |

Note: (*) indicates the best polynomial order for each quantile.

#### D.2 Functional Diversity

Table D7: Quantile regression statistics of the Functional Richness and Fire Return Time relationship for the $N = 10$ Mediterranean simulations (Med-10).

| Quantile | Polynomial Order | AIC | R-squared | Wald test Statistics | Wald test p-value |
|---|---|---|---|---|---|
| 0.85 | 1 | -65509.6 | 0.068 | 1995.4 | 0.0 |
| | 2 (*) | -65951.7 | 0.069 | 1259.5 | 0.0 |
| | 3 | -65747.5 | 0.133 | 15070.8 | 0.0 |
| | 4 | -65729.6 | 0.134 | 18151.2 | 0.0 |
| | 5 | -65496.4 | 0.141 | 28641.3 | 0.0 |
| 0.90 | 1 | -57325.9 | 0.077 | 1797.8 | 0.0 |
| | 2 (*) | -57967.1 | 0.077 | 1757.7 | 0.0 |
| | 3 | -57879.5 | 0.149 | 18613.6 | 0.0 |
| | 4 | -57762.6 | 0.150 | 20963.4 | 0.0 |
| | 5 | -57481.5 | 0.159 | 34657.8 | 0.0 |
| 0.95 | 1 | -41172.8 | 0.103 | 1111.9 | 0.0 |
| | 2 | -39340.9 | 0.106 | 5773.4 | 0.0 |
| | 3 (*) | -41580.6 | 0.181 | 46676.1 | 0.0 |
| | 4 | -41216.8 | 0.183 | 59891.8 | 0.0 |
| | 5 | -41507.2 | 0.190 | 60368.7 | 0.0 |

Note: (*) indicates the best polynomial order for each quantile.

Table D8: Quantile regression statistics of the Functional Divergence and fire return time relationship for the $N = 10$ Mediterranean simulations (Med-10).

| Quantile | Polynomial Order | AIC | R-squared | Wald test Statistics | Wald test p-value |
|---|---|---|---|---|---|
| 0.85 | 1 | -6874.5 | 0.074 | 2946.6 | 0.0 |
| | 2 | -7694.0 | 0.079 | 2245.4 | 0.0 |
| | 3 | -8461.9 | 0.084 | 2276.8 | 0.0 |
| | 4 (*) | -8848.2 | 0.094 | 2532.3 | 0.0 |
| | 5 | -8671.8 | 0.104 | 2812.5 | 0.0 |
| 0.90 | 1 | -3223.8 | 0.074 | 2179.2 | 0.0 |
| | 2 | -3659.5 | 0.076 | 1941.6 | 0.0 |
| | 3 | -4245.1 | 0.077 | 1917.4 | 0.0 |
| | 4 (*) | -4708.3 | 0.090 | 1838.6 | 0.0 |
| | 5 | -4658.0 | 0.100 | 2141.2 | 0.0 |
| 0.95 | 1 | 2043.9 | 0.074 | 852.9 | 0.0 |
| | 2 | 2363.2 | 0.075 | 1084.7 | 0.0 |
| | 3 | 2326.2 | 0.075 | 1170.7 | 0.0 |
| | 4 | 1392.0 | 0.088 | 914.2 | 0.0 |
| | 5 (*) | 1305.6 | 0.099 | 1292.1 | 0.0 |

Note: (*) indicates the best polynomial order for each quantile.

Table D9: Quantile regression statistics of the Functional Richness and Fire Return Time relationship for the $N = 50$ Mediterranean simulations (Med-50).

| Quantile | Polynomial Order | AIC | R-squared | Wald test Statistics | Wald test p-value |
|---|---|---|---|---|---|
| 0.85 | 1 (*) | -106665.4 | 0.101 | 3404.3 | 0.0 |
| | 2 | -106512.3 | 0.101 | 4830.2 | 0.0 |
| | 3 | -105285.5 | 0.147 | 23758.9 | 0.0 |
| | 4 | -105005.7 | 0.153 | 40104.5 | 0.0 |
| | 5 | -105003.6 | 0.153 | 40422.2 | 0.0 |
| 0.90 | 1 | -91578.9 | 0.096 | 2895.4 | 0.0 |
| | 2 | -91132.1 | 0.096 | 5371.7 | 0.0 |
| | 3 | -92882.9 | 0.160 | 34549.5 | 0.0 |
| | 4 (*) | -93478.4 | 0.161 | 39103.9 | 0.0 |
| | 5 | -93063.6 | 0.162 | 38708.7 | 0.0 |
| 0.95 | 1 | -68688.4 | 0.066 | 1414.9 | 0.0 |
| | 2 | -67620.1 | 0.073 | 903.4 | 0.0 |
| | 3 | -69750.7 | 0.155 | 50594.3 | 0.0 |
| | 4 | -68935.0 | 0.158 | 80948.9 | 0.0 |
| | 5 (*) | -69948.1 | 0.165 | 50963.1 | 0.0 |

Note: (*) indicates the best polynomial order for each quantile.

Table D10: Quantile regression statistics of the Functional Divergence and Fire Return Time relationship for the $N = 50$ Mediterranean communities (Med-50).

| Quantile | Polynomial Order | AIC | R-squared | Wald test Statistics | Wald test p-value |
|---|---|---|---|---|---|
| 0.85 | 1 | -29649.2 | 0.145 | 7043.7 | 0.0 |
| | 2 (*) | -29720.7 | 0.146 | 7501.3 | 0.0 |
| | 3 | -29673.5 | 0.146 | 7799.1 | 0.0 |
| | 4 | -29705.7 | 0.146 | 8025.6 | 0.0 |
| | 5 | -29287.4 | 0.151 | 12958.0 | 0.0 |
| 0.90 | 1 (*) | -22311.1 | 0.137 | 3308.1 | 0.0 |
| | 2 | -21541.5 | 0.139 | 3983.1 | 0.0 |
| | 3 | -21635.8 | 0.140 | 4205.9 | 0.0 |
| | 4 | -21642.2 | 0.140 | 4261.0 | 0.0 |
| | 5 | -21764.0 | 0.144 | 11001.0 | 0.0 |
| 0.95 | 1 | -12145.7 | 0.120 | 2661.7 | 0.0 |
| | 2 (*) | -12166.8 | 0.126 | 3156.8 | 0.0 |
| | 3 | -11999.6 | 0.126 | 3225.6 | 0.0 |
| | 4 | -11902.7 | 0.126 | 3434.4 | 0.0 |
| | 5 | -11968.9 | 0.127 | 4279.7 | 0.0 |

Note: (*) indicates the best polynomial order for each quantile.

Table D11: Quantile regression statistics of the Functional Richness and Fire Return Time relationship for the $N = 10$ Boreal communities (Bor-10).

| Quantile | Polynomial Order | AIC | R-squared | Wald test Statistics | Wald test p-value |
|---|---|---|---|---|---|
| 0.85 | 1 | -62858.4 | 0.117 | 3018.7 | 0.0 |
| | 2 | -62374.0 | 0.118 | 5582.1 | 0.0 |
| | 3 (*) | -63265.3 | 0.183 | 20682.2 | 0.0 |
| | 4 | -63098.5 | 0.190 | 38491.9 | 0.0 |
| | 5 | -63078.8 | 0.192 | 39273.3 | 0.0 |
| 0.90 | 1 | -55871.6 | 0.115 | 2530.0 | 0.0 |
| | 2 | -55891.7 | 0.115 | 3593.9 | 0.0 |
| | 3 | -55554.3 | 0.196 | 31150.1 | 0.0 |
| | 4 (*) | -55915.2 | 0.197 | 43125.5 | 0.0 |
| | 5 | -55638.9 | 0.203 | 51312.3 | 0.0 |
| 0.95 | 1 | -43149.4 | 0.097 | 1549.0 | 0.0 |
| | 2 | -43642.6 | 0.101 | 1372.2 | 0.0 |
| | 3 | -43963.1 | 0.202 | 52746.3 | 0.0 |
| | 4 | -43927.5 | 0.202 | 53541.4 | 0.0 |
| | 5 (*) | -44149.9 | 0.209 | 91048.6 | 0.0 |

Note: (*) indicates the best polynomial order for each quantile.

Table D12: Quantile regression statistics of the Functional Divergence and Fire Return Time relationship for the $N = 10$ Boreal communities (Bor-10).

| **Quantile** | **Polynomial Order** | **AIC** | **R-squared** | **Wald test Statistics** | **Wald test p-value** |
|---|---|---|---|---|---|
| 0.85 | 1 | -9735.3 | 0.129 | 5993.7 | 0.0 |
| | 2 | -10757.4 | 0.140 | 4042.9 | 0.0 |
| | 3 | -11618.9 | 0.149 | 4526.1 | 0.0 |
| | 4 | -11598.6 | 0.150 | 4792.5 | 0.0 |
| | 5 (*) | -11682.3 | 0.155 | 4634.4 | 0.0 |
| 0.90 | 1 | -6067.7 | 0.116 | 5351.0 | 0.0 |
| | 2 | -6964.3 | 0.125 | 3745.5 | 0.0 |
| | 3 | -7866.3 | 0.129 | 4421.1 | 0.0 |
| | 4 (*) | -7926.1 | 0.129 | 4551.5 | 0.0 |
| | 5 | -7220.7 | 0.140 | 4300.6 | 0.0 |
| 0.95 | 1 | -765.0 | 0.095 | 1565.8 | 0.0 |
| | 2 (*) | -1439.3 | 0.100 | 1616.6 | 0.0 |
| | 3 | -1124.5 | 0.101 | 1562.7 | 0.0 |
| | 4 | -756.1 | 0.104 | 3153.9 | 0.0 |
| | 5 | -823.4 | 0.129 | 3120.8 | 0.0 |

Note: (*) indicates the best polynomial order for each quantile.

### E Biodiversity indices distributions

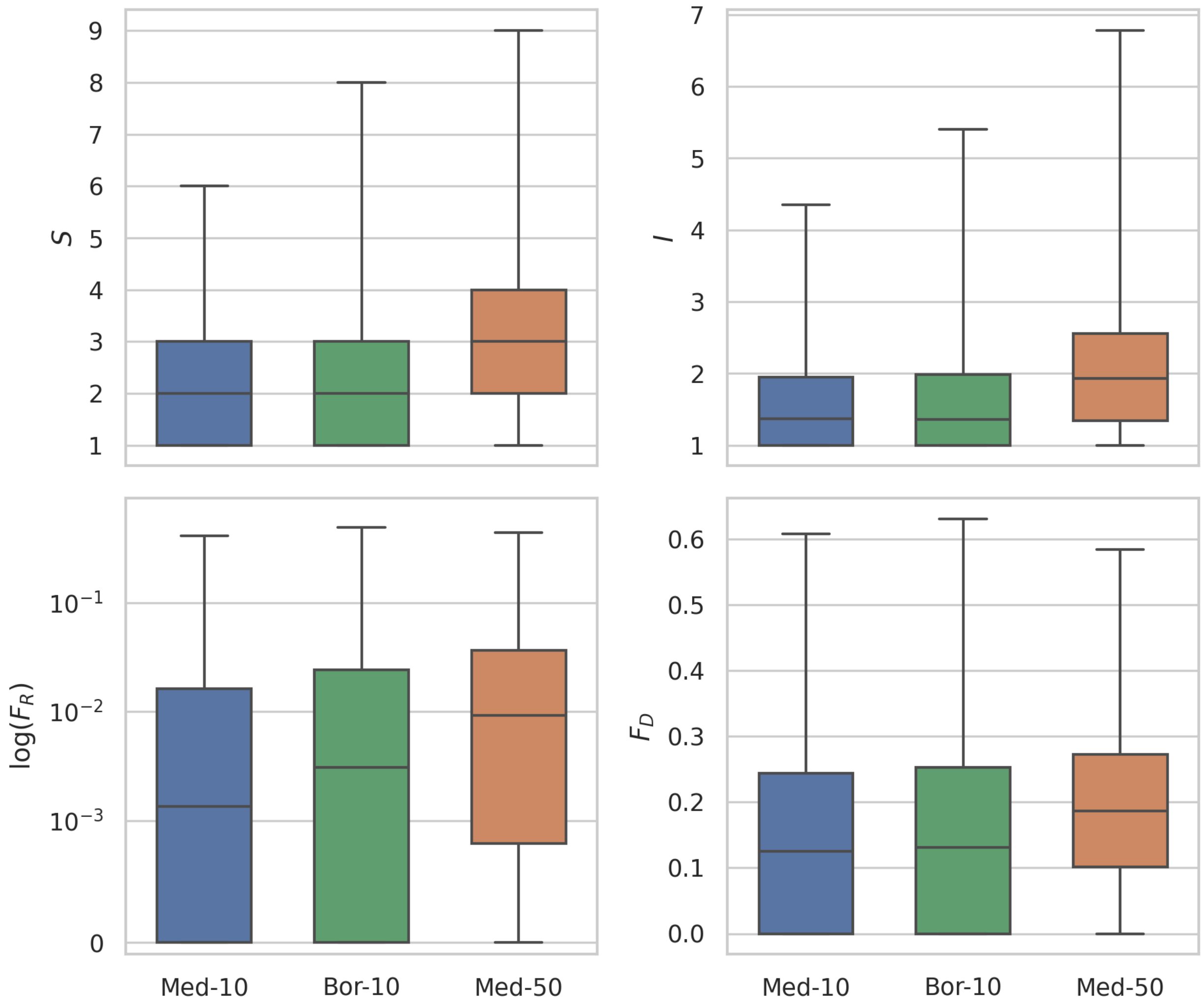


Figure E1: Boxplots of the distributions of biodiversity indicators for the different ecosystem types: Med-10 (blue), Bor-10 (green), Med-50 (orange). $S$ = Species Richness; $I$ = Inverse Simpson Index; $F_R$ = Functional Richness; $F_D$ = Functional Divergence.

### F Biodiversity indices correlations

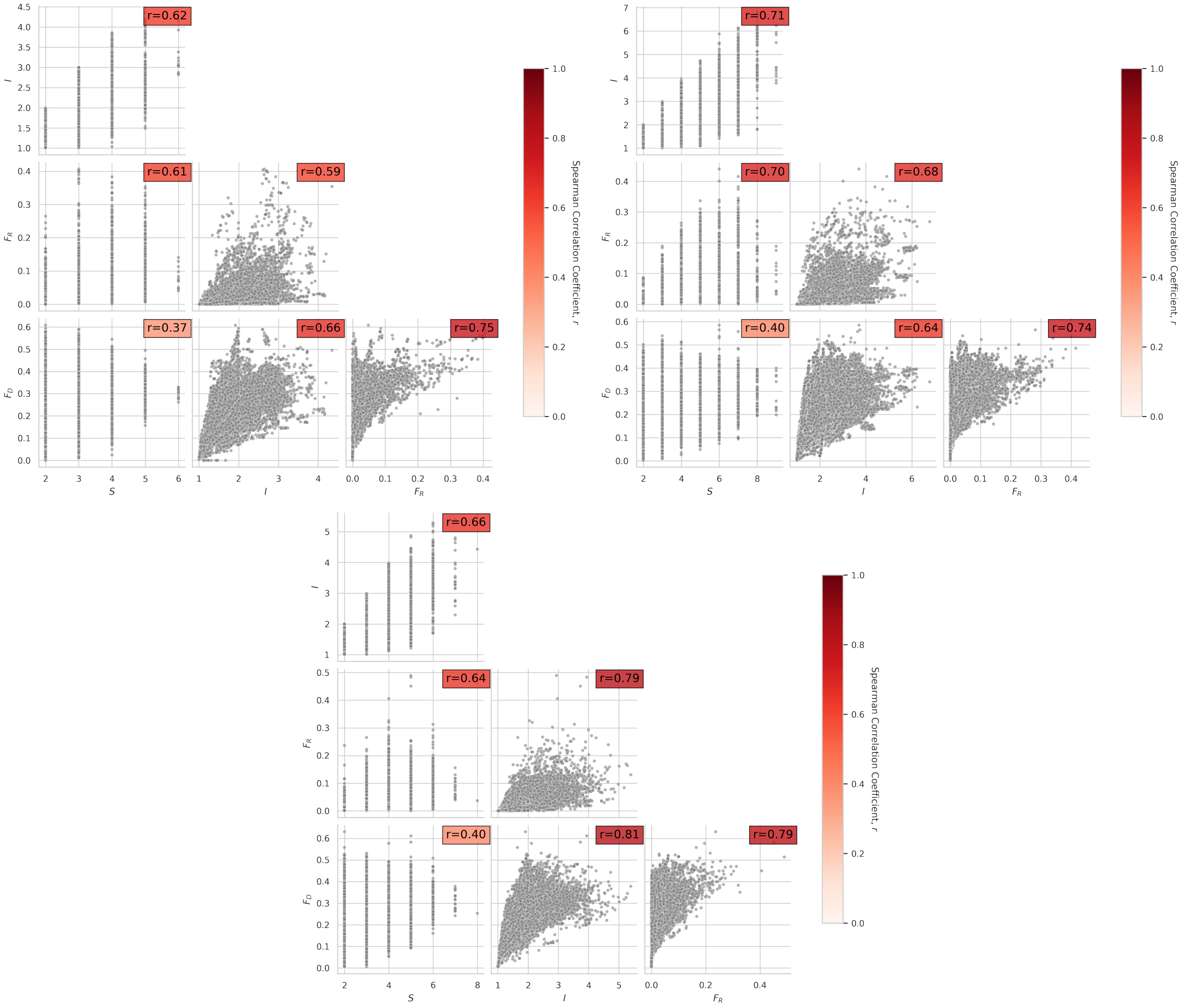


Figure F1: Scatterplots and Spearman's rank-order correlation coefficient for the biodiversity indicators divided by ecosystem type. All correlations resulted significant ($p < 0.001$).

### G Simulations with fixed average fire return time

Here, we apply a series of fixed average fire return times $T_f$ to our model. Specifically, $T_f = 10^x$ yr, where $x$ is extracted from a uniform distribution between 0 and 3, in 61 steps of width 0.05. Using this average return time, the stochasticity of fire events occurrence is simulated using a stationary Poisson process, similarly to the main experiments. The Inverse Simpson Index resulting from the simulations on 300 different Med-10 communities is shown in Fig.G1 bottom-right panel. In this figure, the other eight panels show the results for eight different species communities, thus including all the possible combinations of fixed average return time and initial conditions. Neither single communities nor their aggregation exhibits the hump-shaped relationship found in the main text, where the average fire return time is inversely proportional to the vegetation cover.

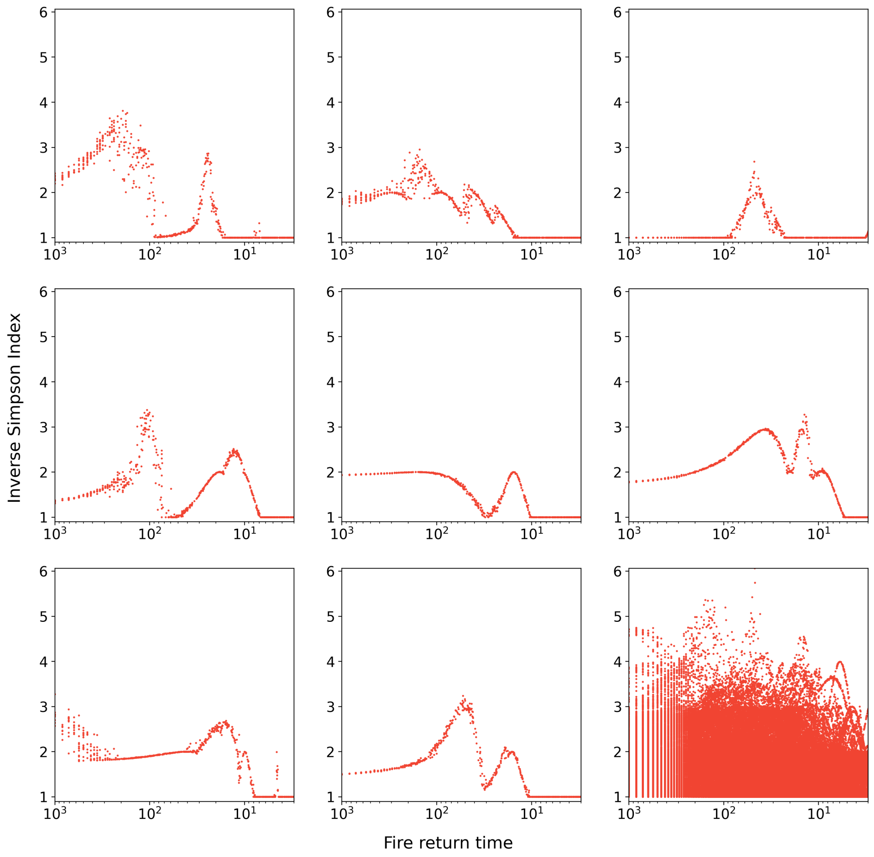


Figure G1: Inverse Simpson Index of communities from 300 simulations of Med-10 with different fixed average fire return time (x-axis). The first eight panels show the results for eight different species communities, while the ninth panel (bottom-right) displays the aggregation of the whole set of 300 communities.

### H Mediterranean simulations including annual grasses

#### *H.1 Extending the range of c values to annual species*

To simulate the annual plants, we extended the range of the colonization rate to larger values up to $C \approx 10$ yr$^{-1}$. To do so, instead of the linear regression used in the main text, we performed an exponential regression to fit the colonization rates obtained from Baudena et al. (2020) as a function of the hierarchy $i$. The regression reads as

$$C(i) = k_1 \exp(k_2 i) + k_3 + \gamma_{C_i}, \tag{H1}$$

where $\gamma_{C_i}$ is a Gaussian distribution with zero mean and $C(i)/3$ standard deviation, and the parameters are $k_1 = 3.734 \times 10^{-4}$yr$^{-1}$, $k_2 = 1.026$, and $k_3 = 4.478 \times 10^{-2}$yr$^{-1}$.

Mortality has been generated as

$$M(i) = \frac{C_{det}(i)}{10} + \gamma_{M_i}, \tag{H2}$$

where $\gamma_{M_i}$ is a Gaussian distribution with zero mean and $M(i)/3$ standard deviation, and $C_{det}(i)$ is the deterministic part of $C(i)$ — that is, $C(i)$ excluding $\gamma_{C_i}$. When generating both $C(i)$ and $M(i)$, we imposed the condition $C(i) > M(i) > 0$.

Fig.H1 shows the colonization and mortality rates generated for $N = 50$ case.

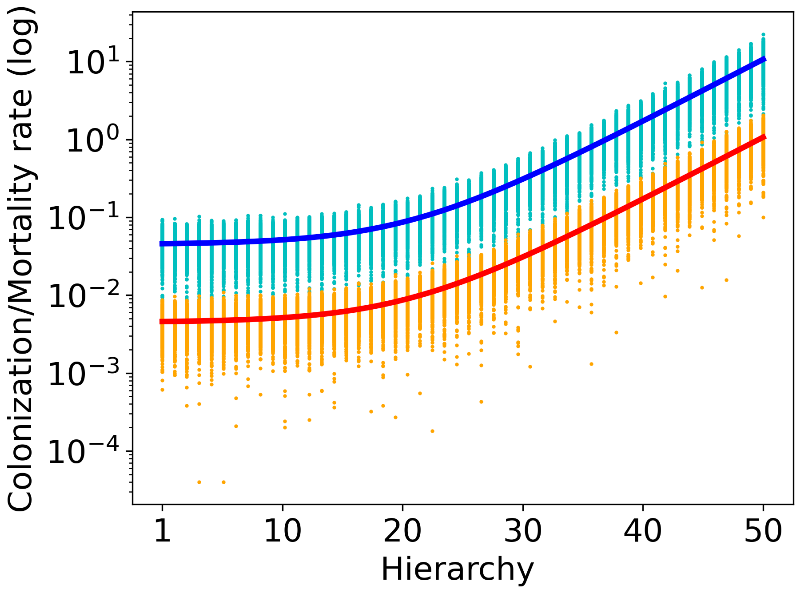


Figure H1: Exponential regression of Baudena et al. (2020) data for colonization (blue line) and mortality (red line) rates. Dots arranged on vertical lines represent the parameters' distribution for 50 species for colonization (cyan dots) and mortality (orange dots).

#### H.2 Compositional and Functional diversity with and without annual species

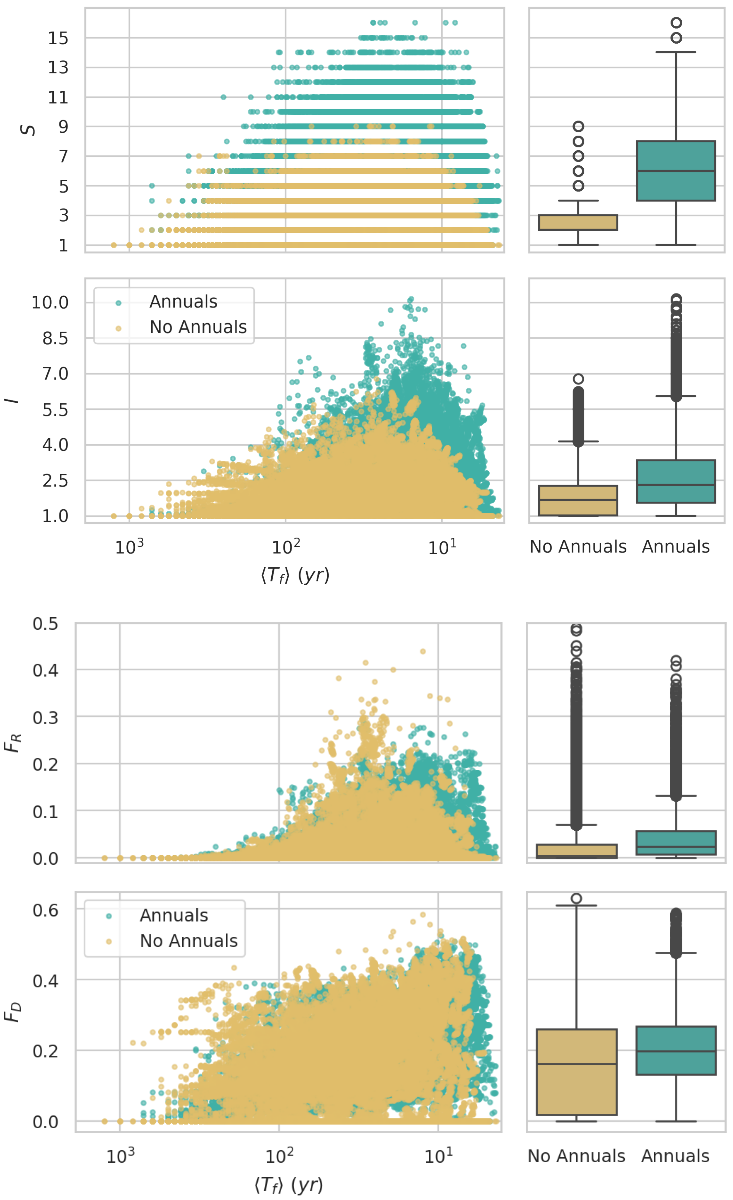


Figure H2: Compositional and Functional diversity for the Mediterranean communities with $N = 50$ initial simulated species generated with (teal) and without (goldenrod) annual plants. Species Richness $S$ (top panels), Inverse Simpson Index $I$ (second-top panels), Functional Richness $F_R$ (second-bottom panels), and Functional Divergence $F_D$ (bottom panels) are plotted as a function of average fire return time $\langle T_f \rangle$ (left panels) and the boxplots of their distributions are plotted for comparison (right panels).

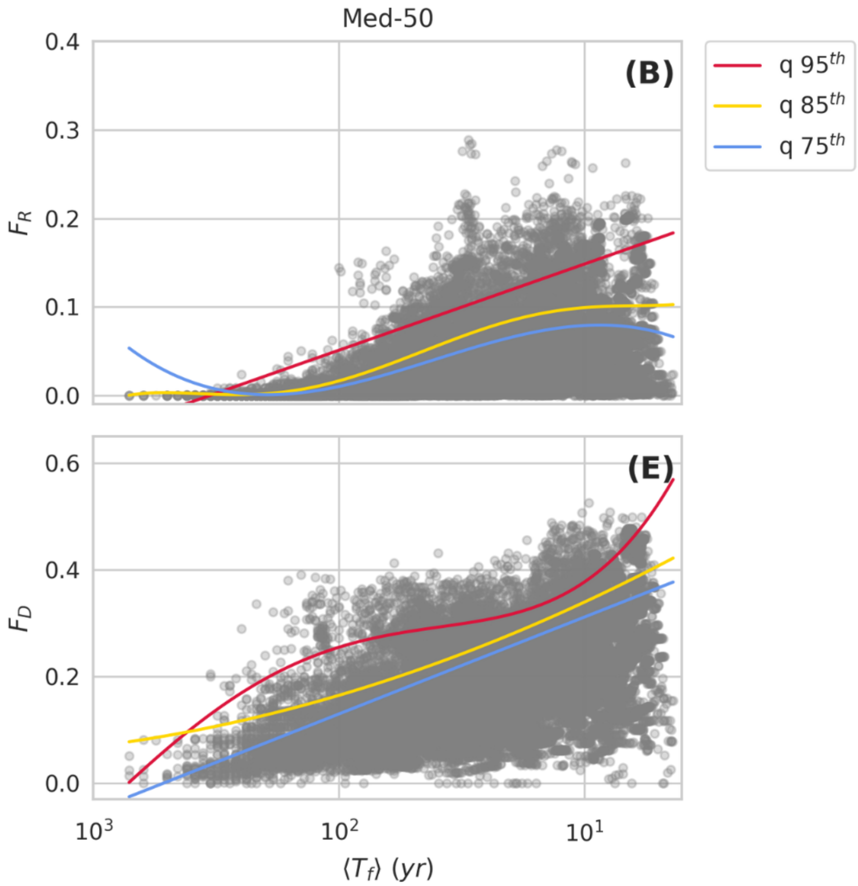


Figure H3: Functional diversity plots over fire frequency for the Mediterranean communities with $N = 50$ initial simulated species generated with annual plants. Functional Richness $F_R$ (top panel) and Functional Divergence $F_D$ (bottom panel) based on the average plant cover are plotted versus average fire return time $\langle T_f \rangle$. The 95$^{th}$ (red), 85$^{th}$ (blue), and 75$^{th}$ (yellow) quantile regressions are reported for each plot, and their polynomial order is chosen based on the lowest AIC value.

#### References Cited Only in Supplementary Information

R. Koenker and S. Portnoy. *Quantile Regression*. ABE, 1996.
